# The ANKS3/BICC1 protein complex is a master post-transcriptional regulator of *NPHP1* ciliopathy-gene transcripts

**DOI:** 10.1101/2024.04.18.588747

**Authors:** Alexia Mahuzier, Gweltas Odye, Valentina Grampa, Albane Bizet, Amandine Viau, Rebecca Ryan, Manon Mehraz, Thierry Blisnick, Euan Clark, Charline Henry, Rémi Piedagnel, Flora Silbermann, Gaelle Hayot, Line De grande, Agathe Kahn, Jean-Marc Plaza, Pauline Krug, Bertrand Knebelmann, Florian Muller, Philippe Bastin, Andreas W. Sailer, Pierre Saint-Mezard, Cécile Jeanpierre, Sigrid Hoffmann, Alexandre Benmerah, Brigitte Lelongt, Marion Delous, Sophie Saunier

## Abstract

Ciliopathies are a class of multi-systemic genetic diseases characterized by ciliary dysfunction. Here, we report a novel *ANKS3* variant in patients with a renal ciliopathy known as nephronophthisis (NPH) associated with hepatic defects. ANKS3 is an ankyrin and sterile alpha motif domain-containing protein that interacts with many NPH proteins as well as with BICC1, an RNA-binding protein involved in renal cystic diseases. The pathogenic effect of the *ANKS3* mutation was validated in the zebrafish mutant and knock-in rat model, the latter showing urine concentration defect and tubular dilatations similar to NPH patients. In addition, cilia morphology and function as well as epithelialization of kidney tubular cells was affected by loss or mutation of *ANKS3*. Finally, our results evidenced that these classically renal ciliopathy-associated phenotypes were linked to the negative regulation of BICC1 by ANKS3 which binds to transcripts of the major NPH gene *NPHP1* and mediates their decay through the AGO2-RISC complex and recruitment into P-bodies. Altogether, our findings suggest that the ANKS3/BICC1 complex is a key post-transcriptional regulator of *NPHP1* transcript stability, providing another level of regulation of cilium biogenesis and kidney homeostasis, as well as an unusual mechanism leading to NPH-related ciliopathies.

## Introduction

Ciliopathies are multi-systemic disorders caused by dysfunction of motile or primary cilia. Whereas motile cilia generate fluid flow or movement of sperm cells, primary cilia act as cellular antennae transmitting extracellular signals essential for tissue development and homeostasis (1, 2). Renal fibrocystic disease, notably nephronophthisis (NPH), is a common manifestation of ciliopathies affecting primary cilia. Characterized by polyuria and polydipsia as a result of urine concentration defect associated to the development of massive renal interstitial fibrosis and cysts at a later stage, NPH is a frequent cause of end-stage renal disease in children and young adults (3). Most proteins encoded by the *NPHP* genes are structural components of different functional subdomains of the cilium such as the transition zone (TZ) and the inversin compartment (InvsC) found at the proximal part of the axoneme, as well as components of the intraflagellar transport (IFT) (3–5). All these proteins play key roles in the biogenesis or function of cilia. The TZ contains the NPHP1, NPHP4 and RPGRIP1L/NPHP8 proteins and forms a molecular filter that selectively controls the transport of ciliary proteins by the IFT machinery. INVS/NPHP2, NPHP3, NEK8/NPHP9 and ANKS6 constitute the InvsC, controlling the establishment of left-right asymmetry and critical ciliary signalling pathways such as flow sensing through polycystins and the canonical Wnt and Wnt/PCP signalling (6, 7). While variants in the TZ genes *NPHP1* and *NPHP4* are mostly associated with juvenile or young adult NPH (8, 9), variants in InvsC genes have been implicated in syndromic forms of infantile NPH with cystic kidneys associated with *situs inversus*, cardiac malformations, and hepatic fibrocystic disease (3). Variants in IFT components (*IFT144/WDR19*, *IFT139/ TTC21B*, *IFT54/TRAF3IP1*, *IFT172/NPHP18*) are found in patients with NPH and retinal and/or bone defects (4, 5, 10, 11). Besides their ciliary functions, NPHP proteins also maintain cell polarity through modulation of cell junctions (NPHP1, NPHP4) and microtubule cytoskeleton dynamics (IFT54, IFT139) (5, 12).

More recently, ANKS3 (a 656 amino acid protein with six ankyrin repeats and a sterile alpha motif [SAM] domain (13)) has been identified as a component of several NPHP modules (14). It interacts with proteins from the TZ (NPHP1, NPHP4, RPGRIP1L) and the InvsC (ANKS6, NEK8, and INVS). These interactions are likely mediated by the ankyrin repeats and the SAM domain, the latter also being involved in the formation of macromolecular complexes with the SAM domains of both ANKS6 and BICC1 (15, 16). BICC1 is an RNA-binding protein that controls the fate of ciliary gene transcripts. Its function relies on its SAM domain that allows polymerization and formation of BICC1 foci, which are modulated by ANKS3 and ANKS6 (15). BICC1 functions with the RISC-associated protein Argonaute-2 (AGO2) system where Argonaute proteins, in association to miRNA, repress translation of transcripts and accumulate them in different RNA processing bodies (17–20). Interestingly, inactivation of BICC1 in mouse, zebrafish and *Drosophila* models as well as *BICC1* heterozygous mutations in humans, result in cystic kidney tubules (17, 21–25). In addition to its role in the control of mRNA expression, BICC1 was recently found to mediate the decay of *Dand5* transcripts in the laterality organs in several species (26, 27). This indicate that in addition to its role as translation regulator, BICC1 can act, in other contexts, as a mediator of mRNA decay for a subset of genes that are still to be characterized.

Here, we report the first case of an *ANKS3* variant in a family with late onset NPH and hepatic fibrosis. We also demonstrate that ANKS3 regulates stability of *NPHP1* transcripts, the gene most commonly mutated in NPH, by modulating its binding to BICC1 and the interaction of BICC1 with the RISC-AGO2. These findings thus identify the BICC1-ANKS3 protein complex as a key regulator of the stability of mRNAs critical for the biogenesis and physiological functions of primary cilia and kidney homeostasis.

## Results

### A homozygous *ANKS3* missense variant causes syndromic nephronophthisis in humans and similar defects in a rat model

Using whole exome sequencing (WES) combined with homozygosity mapping, we identified a homozygous c.806C>T variant in exon 8 of the *ANKS3* gene (NM_133450.3) segregating with the disease in a consanguineous family with three affected siblings exhibiting a late onset NPH (kidney failure after 25 years of age) and hepatic fibrosis (Figure 1A, B, Table I). This very rare variant (Genome Aggregation Database (gnomAD) allele frequency 2x10^-6^), affects an evolutionary conserved amino acid (Table I) and leads to a p.P269L substitution predicted to be deleterious by Polyphen-2 and scale-invariant feature transform (SIFT) algorithms (Table I). This variant sits in the N-terminal region, between the ankyrin repeats and the serine rich region (Figure 1C). No additional biallelic variant could be detected in any other gene encompassed within the 5.9 Mb homozygous candidate region (Supplementary Figure 1A).

**Table I:** Mutations of *ANKS3* in 3 siblings of a consanguineous family with late-onset nephronophthisis and hepatic fibrosis.

| Family Individual | Ethnic origin | Refseq Accession | Nucleotide alteration | Amino Acid Change | PP2 | SIFT | CADD | AF (gnomAD) | zygosity | Amino acid sequence conservation | Kidney phenotype | Extrarenal phenotype |
| --- | --- | --- | --- | --- | --- | --- | --- | --- | --- | --- | --- | --- |
| II.1 | Portugal | NM_133450 | c.806C>T | p.Pro269Leu | 0.998 | 0 | 27.9 | 0.000002053 | Hom | *Danio reno* | NPH; ESKF 31 yo<br>Polyuria-polydypsia<br>Renal ultrasound: Cortico-medullary cysts<br>Mild dedifferentiation<br>Proteinuria 1.60g/24h | none |
| II.2 | Portugal | NM_133450 | c.806C>T | p.Pro269Leu | 0.998 | 0 | 27.9 | 0.000002053 | Hom | *Danio reno* | NPH ; ESKF 25 yo<br>Renal cysts | Biliary cysts |
| II.3 | Portugal | NM_133450 | c.806C>T | p.Pro269Leu | 0.998 | 0 | 27.9 | 0.000002053 | Hom | *Danio reno* | NPH ; CKF 27 yo ; ESKF 33 yo<br>Renal cysts<br>Interstitial fibrosis | Biliary cysts |

*Hom* : homozygous, *NPH* : nephronophthisis, *ESKF*: End-Stage Kidney Failure, *CKF*: Chronic Kidney Failure, *yo*: year old, *PP2*: PolyPhen-2, *CADD*: Combined Annotation Dependent Depletion, *AF*: Allele Frequency.

**Fig. 1.**
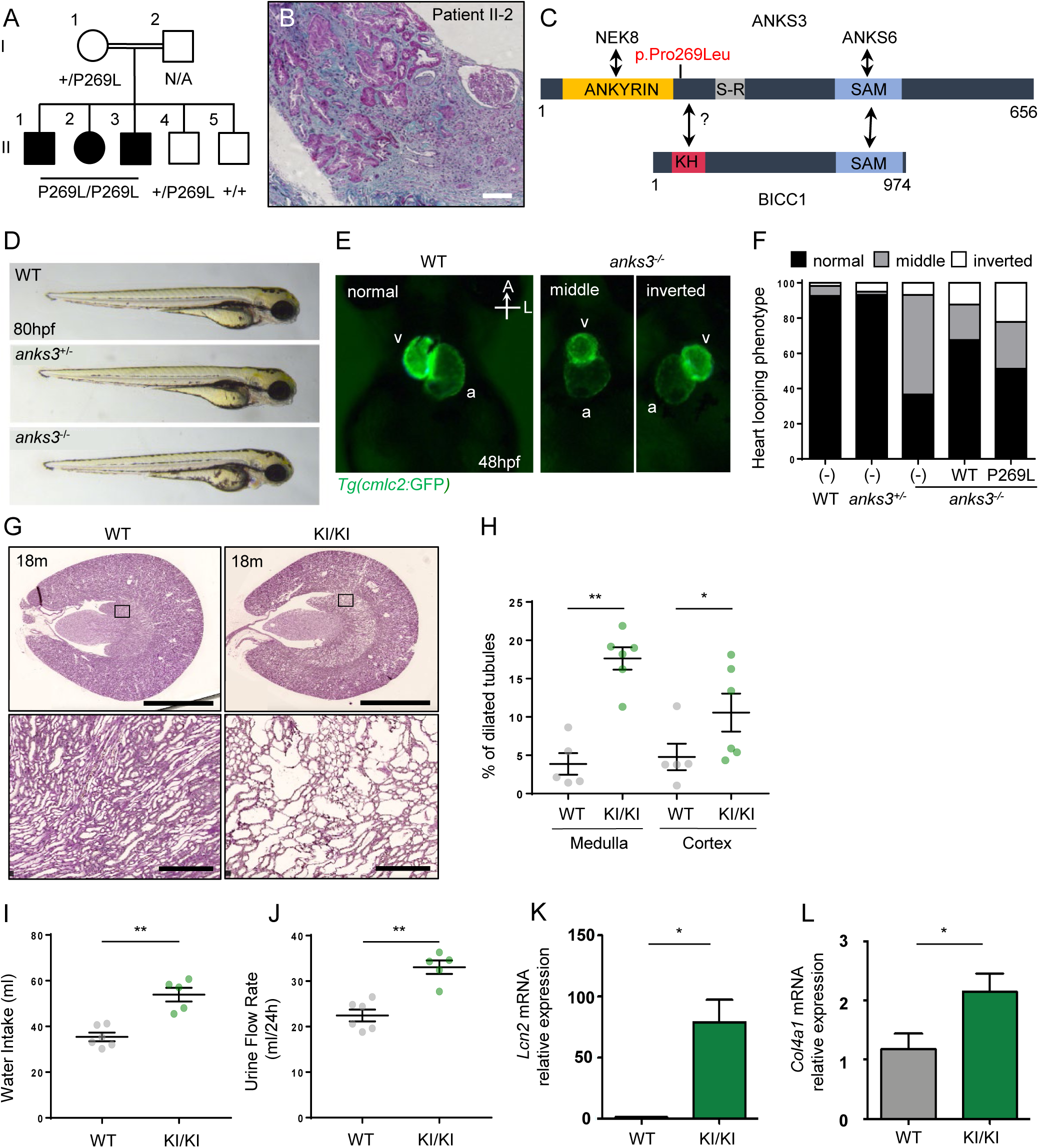
ANKS3 p.P269L mutation causes nephronophthisis-related ciliopathies: **(A)** Pedigree of the consanguineous family with three affected individuals presenting with late onset nephronophthisis (black). **(B)** Hematoxylin and eosin-stained section from a kidney biopsy from patient II.2 showing massive interstitial fibrosis (blue) and focally dilated and atrophic renal tubules. Scale bar, 100 µm. **(C)** Structure of the ANKS3 protein with known BICC1, NEK8 and ANKS6 interaction domains and position of the mutation p.Pro269Leu. **(D)** Gross morphology of WT, heterozygous, and *anks3^ii002^* mutant embryos at 80 hours post-fertilization (hpf). **(E)** Heart looping phenotype of WT and *anks3* mutant embryos at 48 hpf visualized by transgenic line *Tg(cmlc2:GFP)* labelling of the two cardiac chambers (v, ventricle; a, atrium). Ventral views, anterior at top. **(F)** Quantification of heart looping phenotype (%) in WT (n=69), heterozygous (n=126), and *anks3* mutant (n=60) embryos with or without injection of GFP-tagged human WT (n=63) or p.P269L mutated ANKS3 (n=33). **(G-H)** Periodic Acid Schiff-stained section from rat kidney at 18 months from WT and KI/KI (G) showing cystic dilatations in the medullary and cortex regions in the mutant. Scale bar, 5µm (upper panel); 250µm (lower panel). Quantification of percentage of dilated tubules in the medulla and cortex at 18 months for WT (n=5) and KI/KI (n=6) (H). Error bars represent the sem. p-values were determined with Mann-Whitney test. **p≤0.01; *p≤0.05 ns, not significant p>0.05. **(I-J)** Biological data from WT (n=6) and KI/KI (n=6) rats at 18 months showing the water intake (I), the urine flow rate (J) over 24h period. Error bars represent the sem. p-values were determined with Mann-Whitney test. **p≤0.01. **(K-L)** qPCR analyses of relative *Lcn2* (K) and *Col4a1* (L) mRNAs expression normalized to *Hprt* in rat kidney at 18-20 months. Values are the mean ± sem. from 3 animals. *p*-values were determined by Mann-Whitney test. *p≤0.05.

To evaluate the pathogenicity of the identified p.P269L variation, we turned to the zebrafish model, widely used to study ciliopathies. We generated a TALEN-induced *anks3* mutant zebrafish line that presented a frameshift mutation in exon 4 (*ii002*) (Supplementary Figure 1B-D). As previously reported for *anks3* morphants (28), *anks3^-/-^* embryos had normal gross morphology (Figure 1D) but exhibited abnormal heart looping suggestive of defective left-right asymmetry establishment (Figure 1E, F), a well-defined ciliopathy-related phenotype. Injection of human WT *ANKS3* mRNA allowed a partial restoration of the heart looping defects, while the injection of mRNA encoding for the p.P269L variant did not (Figure 1F), thus confirming its pathogenicity.

Then in order to determine the implication of ANKS3 in kidney function, we generated a rat knock-in (KI) model that harbours the same p.P269L mutation as the one identified in the family. First analysis of these KI mutant rats revealed an increase of their body weight at 18 months (Supplementary Figure 2A). Examination of kidneys of 12 months old mutant rats revealed tubular dilatations in the medullary region affecting mainly the collecting ducts (AQP2+), while the other tubular segments appeared without abnormalities (Supplementary Figure 2B, C). At 18 months, the dilatations extended to the cortex region in AQP2 positive tubules, while other dilated tubules were barely stained, suggesting a partial dedifferentiation at this stage (Figure 1G, H; Supplementary Figure 2B). Analysis of kidney function of 18 and 20 months old KI rats using metabolic cages detected a significant increase of water uptake and urine volume flow rate (Figure 1I, J), suggestive of a defect in urine concentration as observed in patients. However, at this stage no elevation of plasma blood urea nitrogen (BUN) was detectable in mutant animals (Supplementary Figure 2D), nevertheless urinary osmolality was significantly decreased in mutant rats at 20 months (Supplementary Figure 2E). At the histological level, kidney fibrosis was not different in mutant and control animals; however, qPCR analysis showed that profibrogenic and tubular damage markers *Col4a1, Tgfβ, Lcn2* and *Kim1* were upregulated in 18-20 months old mutant rat kidneys (Figure 1K, L and Supplementary Figure 2F, G). As two out three patients carrying the p.P269L homozygote variant presented with hepatic defects, we also analysed histological sections of liver of 12 months old mutant rats and detected an increase of hepatic fibrosis (Supplementary Figure 2H). Overall, KI P269L rats developed histological and physiological abnormalities reminiscent of the clinical observations seen in NPH patients.

Altogether, these data identify and validate the p.P269L variant in *ANKS3* as a new cause of NPH.

### ANKS3 controls cellular processes commonly altered in NPH, cell epithelialization and cilium biogenesis

Physiopathological mechanisms leading to NPH include a wide range of cellular processes, of which cell epithelialization and cilium biogenesis are likely to be the most important (5, 12, 29). To explore the role of ANKS3 in these processes, we established a stable short hairpin RNA (shRNA)-mediated knock-down of *Anks3* in murine Inner Medullar Collecting Duct (mIMCD3) cells (KD_ANKS3), which resulted in a 80% reduction of the protein level (Supplementary Figure 3A). Stable expression of either GFP alone (hereafter, KD_GFP) or human WT or p.P269L ANKS3-GFP fusions (hereafter, KD_WT or KD_P269L, respectively) in KD_ANKS3 cells was subsequently performed by lentiviral infection (Supplementary Figure 3A).

As polarity and epithelialization defects in renal tubular cells have been associated with alteration of NPH genes, we first investigated the potential role of ANKS3 in these processes on confluent monolayers of mIMCD3 cells grown on filters. Immunofluorescence analyses revealed that the apical marker GP135 was not detected at the apical membrane in both KD_GFP and KD_P269L cells (Figure 2A) and that cell height was decreased (Figure 2B). Moreover, we performed a calcium-switch assay and measured the trans-epithelial resistance (TER) during re-formation of tight junctions. KD_GFP and KD_P269L cells showed a delay in tight junction formation as monitored by decreased TER compared to control and KD_WT cells (Figure 2C). These phenotypes thus indicated that KD_GFP and KD_P269L cells failed to properly polarize.

**Fig. 2.**
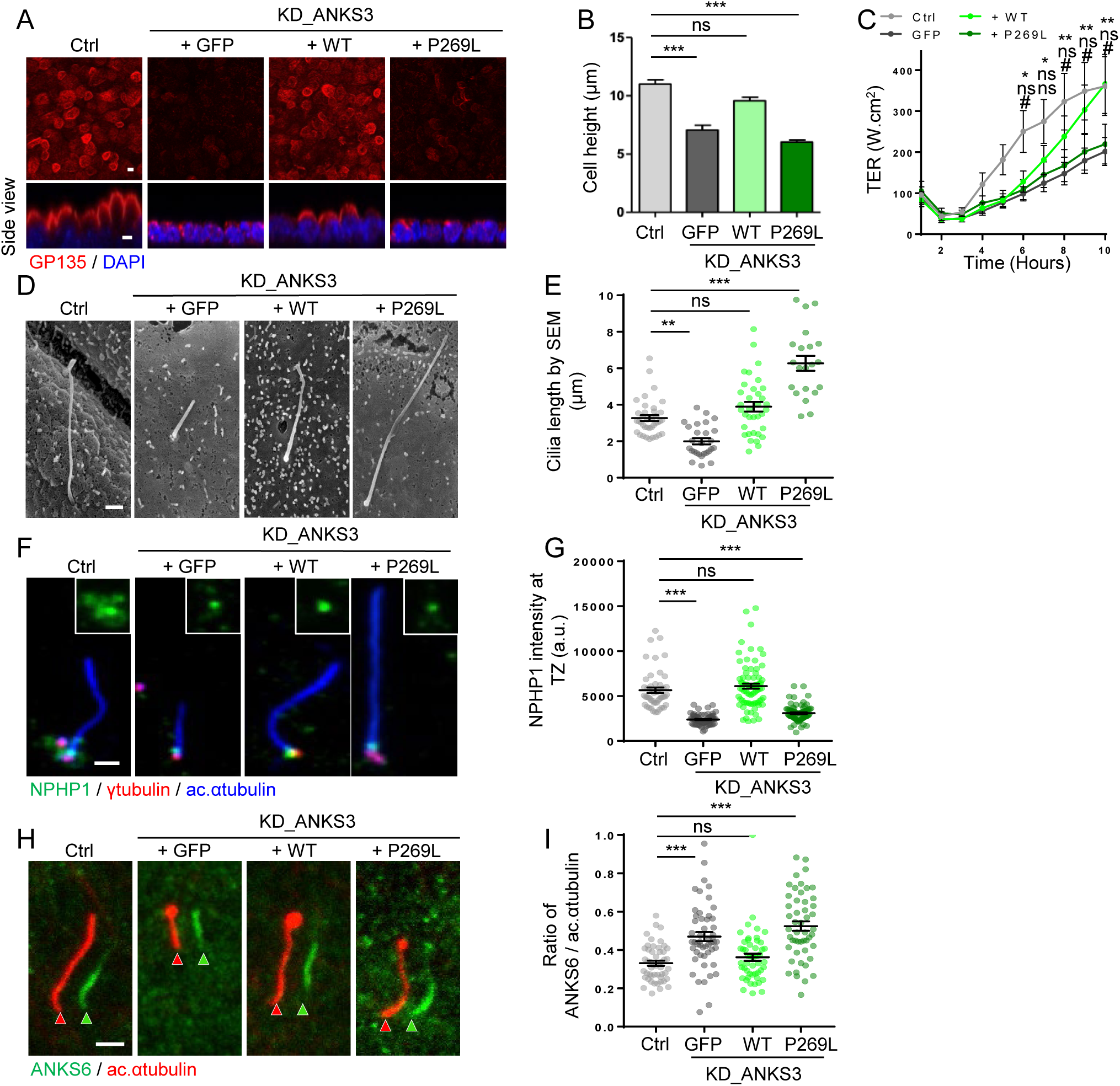
ANKS3 loss of function and NPH-related p.P269L mutation leads to ciliary and polarity defects: **(A-B)** Subcellular localization of the apical marker GP135 (A, red) in Ctrl, KD_GFP, KD_WT, and KD_P269L mIMCD3 cells grown on filter. Scale bar, 10 µm. Quantification of the cell height (B) is based on 3 independent experiments. n>20 cells. **(C)** Measurement of transepithelial resistance (TER) of Ctrl, KD_GFP, KD_WT, and KD_P269L mIMCD3 cells at different time points after Ca^2+^ switch assay of 3 independent experiments; *comparison of Ctrl vs. KD_GFP; # comparison of Ctrl vs. KD_P269L. **(D-E)** Scanning electron microscopy analysis of primary cilia (D) in Ctrl and KD_GFP, KD_WT, and KD_P269L mIMCD3 cells. Scale bar, 1µm. Quantification of cilia length (E) is based on 3 independent experiments. n>25 cells. **(F-G)** Immunostaining of NPHP1 (green), g-tubulin (red) and acetylated α-tubulin (blue) in Ctrl and KD_GFP, KD_WT, and KD_P269L mIMCD3 cells (F). Scale bar, 1µm. Quantification of NPHP1 fluorescence intensity at the TZ (G) is based in 3 independent experiments. **(H-I)** Ciliary distribution of ANKS6 in mIMCD3 cells stained for ANKS6 (green) and acetylated α-tubulin (red) (H). Scale bar, 1µm. Quantification of the size of the InvsC (I) is based on 3 independent experiments. in KI/KI and WT rats. (B, C, E, G, I) Values are the mean ± sem. *p*-values were determined by one-way ANOVA followed by Dunn’s multiple comparison test; ns: non-significant *p* > 0.05; \**p*≤0.05; \*\**p* ≤0.01; ***p≤0.001.

We next analysed ciliogenesis by scanning electron microscopy (SEM, Figure 2D, E) and immunofluorescence (Figure 2F-I) and showed that the loss of ANKS3 (KD_GFP) resulted in shorter cilia while re-expressing WT ANKS3 rescue cilia shortening. Strikingly, the expression of the P269L variant led to longer cilia (Figure 2D-F and Supplementary Figure 3B), without altering the percentage of ciliated cells in both mIMCD3 cells and fibroblasts derived from patient (Supplementary Figure 3C, D). The impact of P269L variant on increased cilia length was confirmed in KI rat kidney sections and in patient fibroblasts, as well as in the kidney biopsy of one of the NPH patients (Supplementary Figure 3E-I). Of note, such elongation of cilia was previously observed in NPH conditions, including *NPHP1* (30), *CEP290* (31), *CEP83* (32) and *IFT54/TRAF3IP1* (5).

The impact of ANKS3 (KD or pathological conditions) on the integrity of the NPH-related ciliary subdomains was analysed. We first focused on TZ proteins. Interestingly, we observed a decreased staining of both NPHP1 and TMEM67 at TZ in KD_ANKS3 cells (Figure 2F, G and Supplementary Figure 3J, K). In contrast, in KD_P269L cells, only NPHP1 levels were decreased (Figure 2F, G and Supplementary Figure 3J, K). Finally, the amount of NPHP4 was not modified in all conditions indicating that not all TZ components are affected (Supplementary Figure 3L). We then investigated the localisation of InvsC components. As expected, NEK8, ANKS6 and INVS were found at the proximal half of the axoneme in control mIMCD3 cells, WT rat kidney tubular epithelial cells and control fibroblasts (Figure 2H, I and Supplementary Figure 4A-H). In contrast, in all P269L cells, a drastic extension of the InvsC was observed with staining of InvsC components observed all along the axoneme (ANKS6, Figure 2H, I and Supplementary Figure 4C-F; NEK8, Supplementary Figure 4A, B; INVS, Supplementary Figure 4G, H). A relative InvsC extension was also noticed in KD_GFP cells in which InvsC components were found all along the axoneme of the observed shorter cilia (Figure 2H, I; Supplementary Figure 4A, B, G, H). Nevertheless, in the latter case, it is unclear whether the observed shorter cilia were restricted to InvsC only or if InvsC proteins were mislocalized all along the cilium, as observed in the elongated cilia of KD_P269L.

As these ciliary compartmentalization defects were likely associated with an alteration of ciliary composition and function, we next examined the localization of key ciliary signaling proteins known to be affected in NPH conditions. We observed that the ciliary translocation of Smoothened (Smo), a typical feature of the Hedgehog pathway (Hh) activation upon agonist (SAG) treatment, was decreased in both KD_GFP and KD_P269L cells compared to controls (Supplementary Figure 4I, J). In agreement with a defective response to Hh pathway activation in *ANKS3* conditions, expression of *Patched1* showed a 50% decrease in response to SAG in KD_GFP and KD_P269L cells (Supplementary Figure 4K) as well as in fibroblasts from the p.P269L patient (Supplementary Figure 4L). In addition, polycystin-2 (PC2), required for cilia-dependent mechanosensation, which was found at the proximal part of the ciliary axoneme in control mIMCD3 cells, was absent from the cilium of KD_GFP cells while it was distributed all along the axoneme in KD_P269L cells (Supplementary Figure 4M, N). Alteration of the cilium function was also observed *in vivo* in *anks3* zebrafish mutants: a decreased beating of cilia in the Kuppfer’s vesicle was associated to the alteration of cilium-dependent calcium signalling (Supplementary Figure 1E-H).

Altogether, these results evidenced that ANKS3, similarly as other NPH proteins, is important for proper epithelial cell polarity as well as proper cilium length, organization and composition, thus guaranteeing the cilia-dependent signalling pathways.

### Ciliogenesis and epithelial defects observed in KD and mutant ANKS3 conditions are mediated by BICC1

To have a first clue by which mechanism ANKS3 is involved in both cilium biogenesis and epithelial polarity, we studied its subcellular localization. Contrary to fibroblasts where ANKS3 was detected at the basal body (Supplementary Figure 3G), endogenous ANKS3 and the WT ANKS3-GFP fusion localized within granular cytoplasmic structures in polarized mIMCD3 cells and in rat renal tubular cells (Figure 3A, B). The P269L mutation did not alter ANKS3 localization in any of our models (Figure 3A, B; Supplementary Figure 3G). Of note, these staining patterns were consistent with previous data showing the localization of overexpressed ANKS3 at both basal body region and in small aggregates in multi-ciliated cells in the epidermis of *Xenopus laevis* embryos (14) or in cytoplasmic foci when co-expressed with BICC1 in HeLa cells (15).

**Fig. 3.**
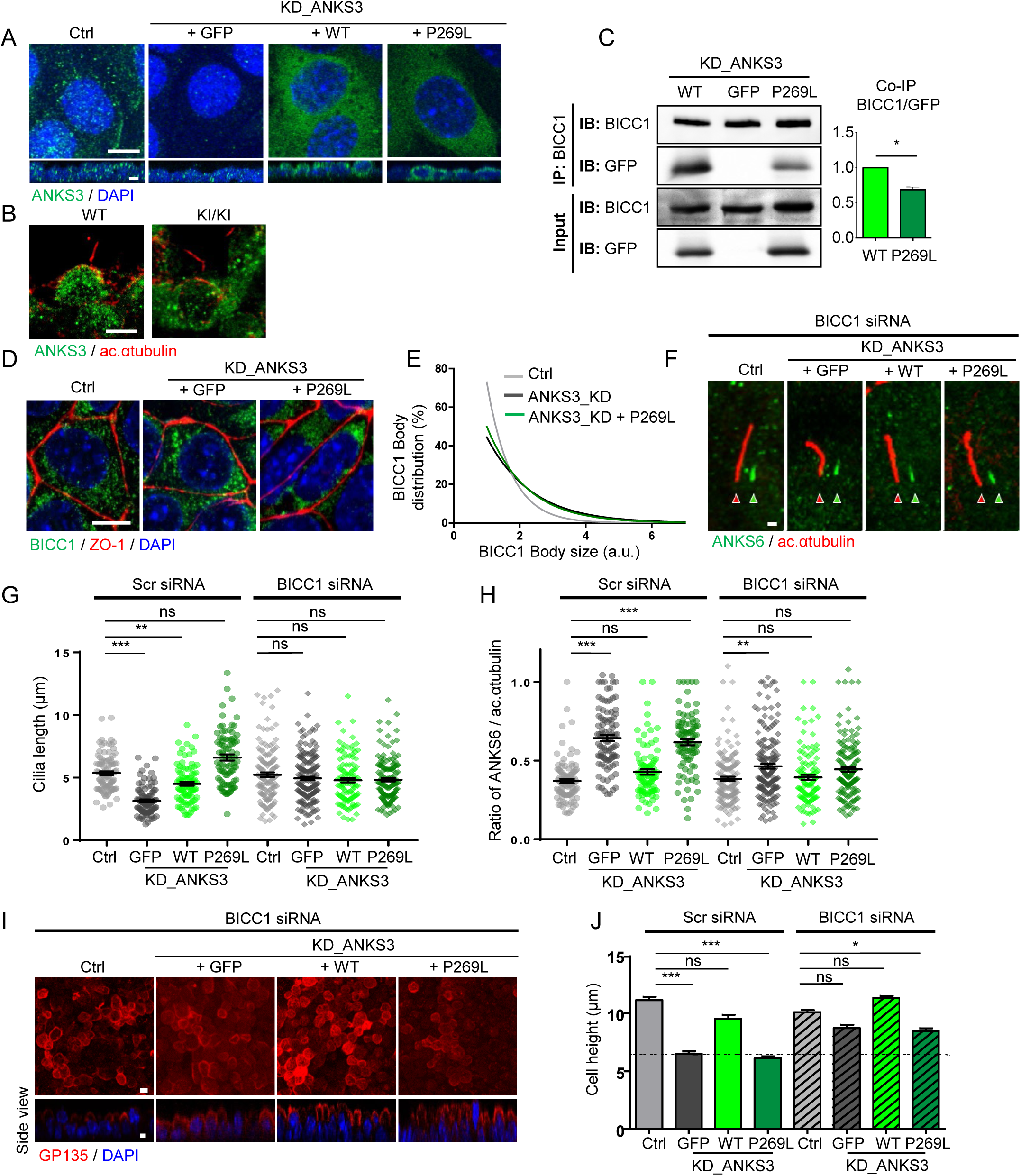
NPH-related p.P269L mutation abrogates ANKS3 interaction with BICC1 and silencing of BICC1 rescues ANKS3 KD-associated defects: **(A-B)** Immunofluorescence analysis of subcellular localization of ANKS3 (green). (A) in control, KD_ANKS3, WT, and mutant mIMCD3 cells. Cells were grown until confluence on glass slides. Scale bars, 10 µm. (B) in rat kidney cross sections at 18 months in WT and KI/KI. Scale bar, 4 µm. **(C)** Effect of p.P269L mutation on interaction between ANKS3 and BICC1. Lysates from mIMCD3 cells with knockdown of ANKS3 (KD_ANKS3) and re-expressing WT ANKS3-GFP, P269L ANKS3-GFP, or GFP alone were immunoprecipitated with an anti-BICC1 antibody. Co-immunoprecipitation of GFP-ANKS3 and BICC1 was quantified by western blot (IB) using GFP and BICC1 antibodies. Error bars represent the sem of 3 independent experiments; *p*-values were determined with the Mann-Whitney test, *p≤0.05. **(D-E)** BICC1 bodies (green) and cell junction (ZO-1, red) in Ctrl and KD_ANKS3 in mIMCD3 cells (D). Scale bars, 5 µm. Quantification of the size of BICC1 bodies (E) using ImageJ spot detector protocol from 3 independent experiments. **(F-H)** Immunofluorescence (F) and quantification of the ciliary length (G) and the ciliary distribution of ANKS6 (H) in Ctrl, KD_GFP, KD_WT, and KD_P269L mIMCD3 cells transfected with scramble or BICC1-targeting siRNA (F, ANKS6, green; acetylated α-tubulin, red). Scale bar, 1µm. ANKS6 distribution (H) was quantified as the ratio of ANKS6 staining length and total cilia length (acetylated α-tubulin staining). Values are the mean ± sem. of n>100 cells from 3 independent experiments; *p*-values were determined by one-way ANOVA followed by Dunn’s multiple comparison test; \*\*\**p* ≤ 0.001; **p≤0.01; ns, not significant p> 0.05. **(I-J)** Subcellular localization of the apical marker GP135 (red) in Ctrl, KD_GFP, KD_WT, and KD_P269L mIMCD3 cells transfected with BICC1-targeting siRNA and grown on filter prior to 6 hours incubation in standard medium after Ca^2+^ switch assay. Scale bar, 10 µm. Quantification of the cell height (J) is based on 3 independent experiments. Values are the mean ± sem. of n>20 cells. *p*-values were determined by one-way ANOVA followed by Dunn’s multiple comparison test; \*\*\**p* ≤ 0.001; *p<0.05; ns, not significant *p*> 0.05.

Our observations prompted us to explore the impact of the P269L variant on the functions of the ANKS3/BICC1 complex. Co-immunoprecipitation experiments showed that the P269L variant interacted less with BICC1 than WT ANKS3 (Figure 3C). ANKS3 was shown to control BICC1 polymerization and the number/size of BICC1 bodies (15). As expected, we observed that the size of BICC1 bodies was increased both in KD_ANKS3 mIMCD3 cells (Figure 3D, E) and patient fibroblasts (Supplementary Figure 5A). Then, to test whether ANKS3-associated phenotypes were linked to impaired BICC1 function, we modulated BICC1 expression in ANKS3 KD cell lines using *Bicc1*-targeting siRNA (Supplementary Figure 5B). In these conditions, ciliary defects, i.e. cilium length (short or long, Figure 3F, G) and InvsC biogenesis (Figure 3F, H and Supplementary Figure 5C, D) as well as the polarity phenotypes (GP135 localisation, cell height, Figure 3I, J) were partially or totally rescued by *Bicc1* knockdown.

Altogether, these results confirmed that ANKS3 regulates BICC1 body dynamics and that BICC1 is involved in the mechanisms by which loss of ANKS3 leads to cilia- and non-cilia-related phenotypes.

### ANKS3 functions with BICC1 to regulate *Nphp1* transcript stability

BICC1 is known to regulate the translation of a specific set of mRNA associated with kidney cystogenesis (33) or to mediate the decay of *Dand5* mRNA in the context of laterality organ (26, 27). We hypothesized that the ANKS3-associated phenotypes might be related to its potential involvement in the regulation of BICC1-mediated control of ciliopathy-related transcripts. Most of the ciliary and cell polarity phenotypes observed in ANKS3 conditions were reminiscent to what is observed in NPHP1 loss of function (12, Supplementary Figure 6A-D). Notably, we observed a global decreased expression of NPHP1 in KD_GFP and KD_P269L both by immunofluorescence, at the TZ (Figure 2F, G) and in the cytoplasm (Figure 4A), and by Western blot (Figure 4B). We thus explored the possibility that the downregulation of NPHP1 protein expression was linked to BICC1-mediated regulation of *Nphp1* mRNAs. First, we performed RNA co-immunoprecipitation (RIP) assays and found that *Nphp1* mRNAs were indeed co-immunoprecipitated with BICC1. Interestingly, *Nphp1* transcripts were more abundant in BICC1 RIPs in the absence of ANKS3, or in the presence of the P269L variant, than in control conditions (Figure 4C). This increased interaction was specific to *Nphp1* mRNA since it was not observed for *Adcy6* transcripts (Figure 4C), a known target of BICC1 in kidney epithelial cells (23). Concomitantly, we observed that the total levels of *Nphp1* transcripts were decreased in KD and mutant ANKS3 conditions (Figure 4C, see inputs), and confirmed that this *Nphp1* mRNA decrease was dependent on BICC1 expression (Figure 4D). This suggests that BICC1 control *Nphp1* mRNA level, a function that could be linked to either mRNA degradation/stability or/and transcriptional silencing. We used actinomycin D treatment to block RNA transcription and followed mRNA stability over time. The half-life of *Nphp1* mRNA was decreased in KD_GFP and KD_P269L (Figure 4E), thus supporting a decreased stability of *Nphp1* transcript rather than a decreased transcription. Finally, siRNA-mediated BICC1 depletion rescued shortened *Nphp1* transcript half-life in KD_GFP or KD_P269L cells (Figure 4F), in agreement with the observed restoration of NPHP1 expression at both mRNA and protein levels (Figure 4B, D). Altogether, those results showed that in kidney epithelial cells, ANKS3 acts as a negative regulator of BICC1, which mediates *Nphp11* transcript decay in agreement with results on *Dand5* in the context of the laterality organ (26, 27).

**Fig. 4.**
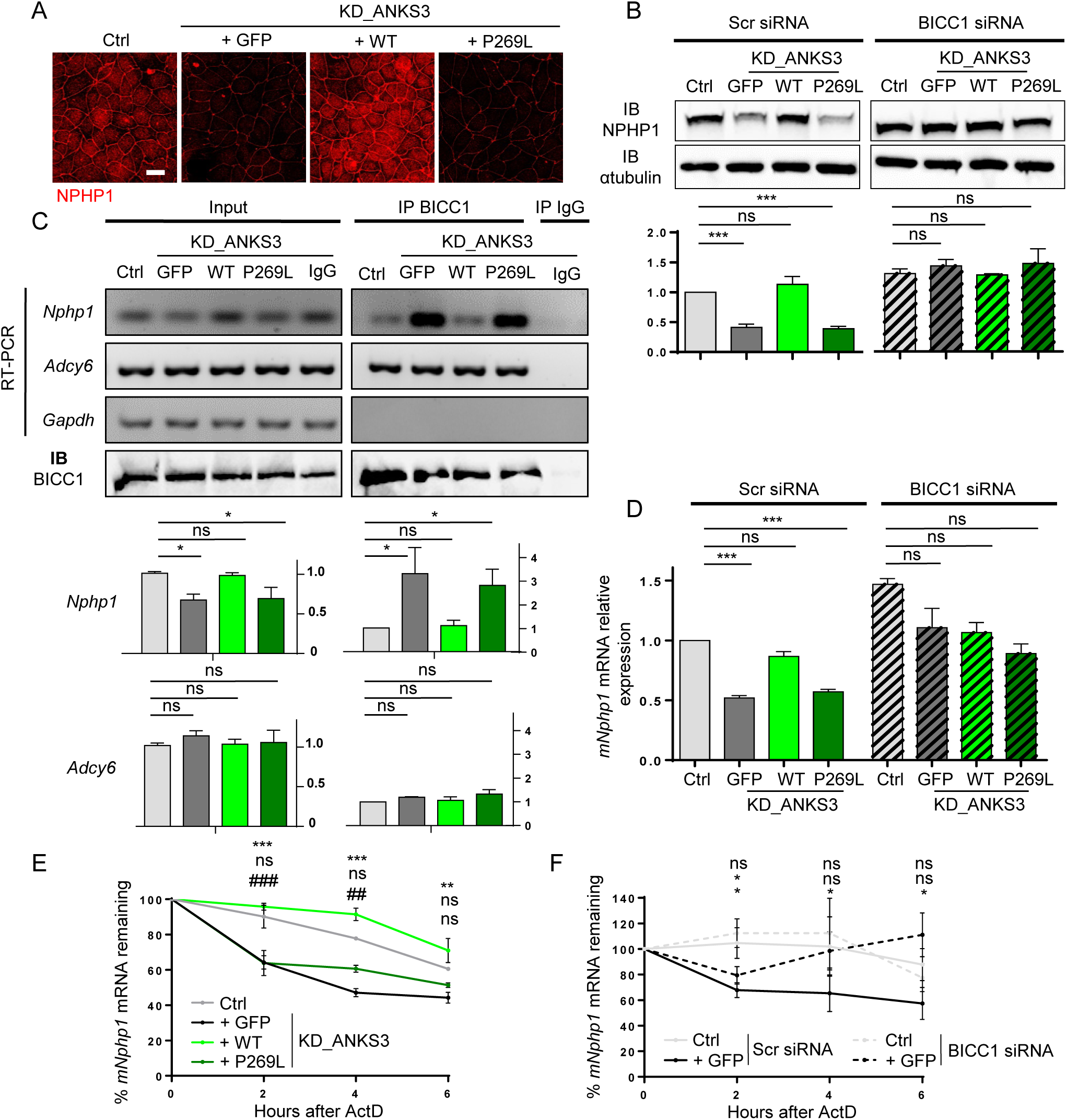
*Nphp1* transcript stability is dependent of BICC1 expression: **(A)** NPHP1 immunostaining in Ctrl, KD_GFP, KD_WT, and KD_P269L mIMCD3 cells. Scale bar, 10µm. **(B)** Western blot analysis of NPHP1 expression in KD_GFP, KD_WT, and KD_P269L mIMCD3 cells transfected with a scramble siRNA or BICC1-targeting siRNA. Quantification of NPHP1 expression by western blot is based on 3 independent experiments. Error bars represent the sem. *p*-values were determined by one-way ANOVA followed by Dunn’s multiple comparison test, ***p≤0.001; ns, not significant *p*> 0.05. **(C)** RNA coimmunoprecipitation of *Nphp1* mRNA with BICC1 in Ctrl, KD_GFP, KD_WT, and KD_P269L mIMCD3 cells. Immunoprecipitates were subjected to RT-PCR for *Nphp1*, *Adcy6* as negative control and *Gapdh*. Western blot with BICC1 antibody (bottom panel) was performed on immunoprecipitates. Quantifications of total and immunoprecipitated *Nphp1* and *Adcy6* mRNAs are based on 4 independent experiments. Values are the mean ± sem. *p*-values were determined by one-way ANOVA followed by Dunn’s multiple comparison test; \**p* ≤ 0.05; ns, not significant p> 0.05. **(D)** qPCR analysis of relative *Nphp1* mRNA expression normalized to *Hprt* in Ctrl, KD_GFP, KD_WT, and KD_P269L mIMCD3 cells transfected with a scramble siRNA or BICC1-targeting siRNA. Values are the mean ± sem. from 3 independent experiments; *p*-values were determined by one-way ANOVA followed by Dunn’s multiple comparison test; \*\*\**p* ≤ 0.001; ns, not significant p> 0.05. **(E-F)** qPCR analysis of murine *Nphp1* mRNA decay kinetics after actinomycin D (ActD) treatment in Ctrl, KD_GFP, KD_WT, and KD_P269L mIMCD3 cells (E), and transfected with scramble or BICC1-targeting siRNA (F). Values are the mean ± sem. of 3 independent experiments; *comparison of Ctrl vs. KD_GFP; # comparison of Ctrl vs KD_P269L; *p*-values were determined by one-way ANOVA followed by Dunn’s multiple comparison test; ns: non-significant *p*>0.05, **/## p ≤ 0.01, ***/### p ≤0.001.

**Fig. 5.**
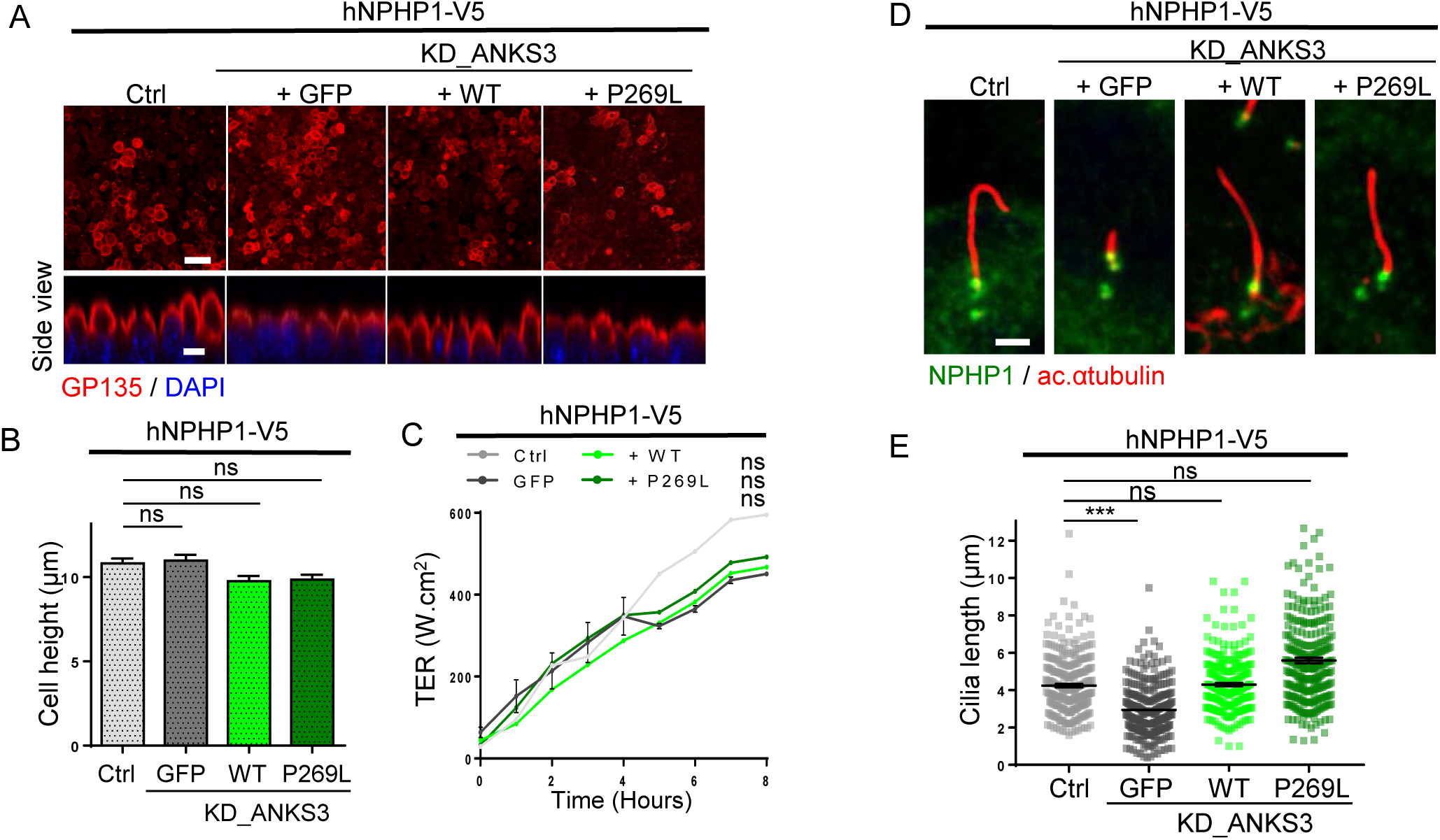
Reexpression of NPHP1 rescues epithelialisation and ciliogenesis defects in KD_ANKS3 condition, but not cilia length of the KD_GFP: **(A-C)** Immunofluorescence of the apical marker GP135 (A, red) in Ctrl, KD_GFP, KD_WT, and KD_P269L mIMCD3 cells stably expressing human NPHP1-V5 and grown for 6 h in standard medium after Ca^2+^ switch assay. Scale bar, 10 µm. Quantification of the cell height (B) is based on 3 independent experiments. Values are the mean ± sem. of n>20 cells. *p*-values were determined by one-way ANOVA followed by Dunn’s multiple comparison test. ns: non-significant p> 0.05. (**C**) Measurement of transepithelial resistance (TER) of Ctrl, KD_GFP, KD_WT, and KD_P269L mIMCD3 cells stably expressing human NPHP1-V5 at different time points after Ca^2+^ switch assay. Values are the mean ± sem. of 3 independent experiments; *p*-values were determined by one-way ANOVA followed by Dunn’s multiple comparison test; ns: non-significant, p>0.05. **(D, E)** Immunofluorescence of NPHP1 (green) and acetylated γ-tubulin (red) in Ctrl, KD_GFP, KD_WT, and KD_P269L mIMCD3 cells stably expressing human NPHP1-V5. Scale bar, 2 µm. Quantification of cilia length is based on 3 independent experiments. Values are the mean ± sem. of n>100 cells. *p*-values were determined by one-way ANOVA followed by Dunn’s multiple comparison test; ns: non-significant *p*>0.05, ***p≤0.001.

To directly investigate the specific contribution of the downregulation of *Nphp1* in the ANKS3-related phenotypes, we performed rescue experiments through stable expression of human V5-tagged human NPHP1 (Supplementary Figure 6E). Notably, re-expression of NPHP1 was able to fully rescue the polarity defects observed in KD_GFP and KD_P269L including GP135 expression, cell height and trans-epithelial resistance (Figure 6A-C). These latter results suggest a main contribution of *Nphp1* downregulation in the epithelialization defects observed in ANKS3 conditions. Moreover, re-expression of NPHP1 also corrected the longer cilia phenotype in KD_P269L (Figure 6D, E), however it did not rescue shorter cilia, InvsC elongation (Figure 6D, E and Supplementary Figure 6F) observed in KD_GFP and KD_P269L cells. These results indicate that *Nphp1* down-regulation does not account for all ciliary phenotypes observed in ANKS3 deficient cells and that probably other transcripts regulated by BICC1/ANKS3 are involved.

**Fig. 6.**
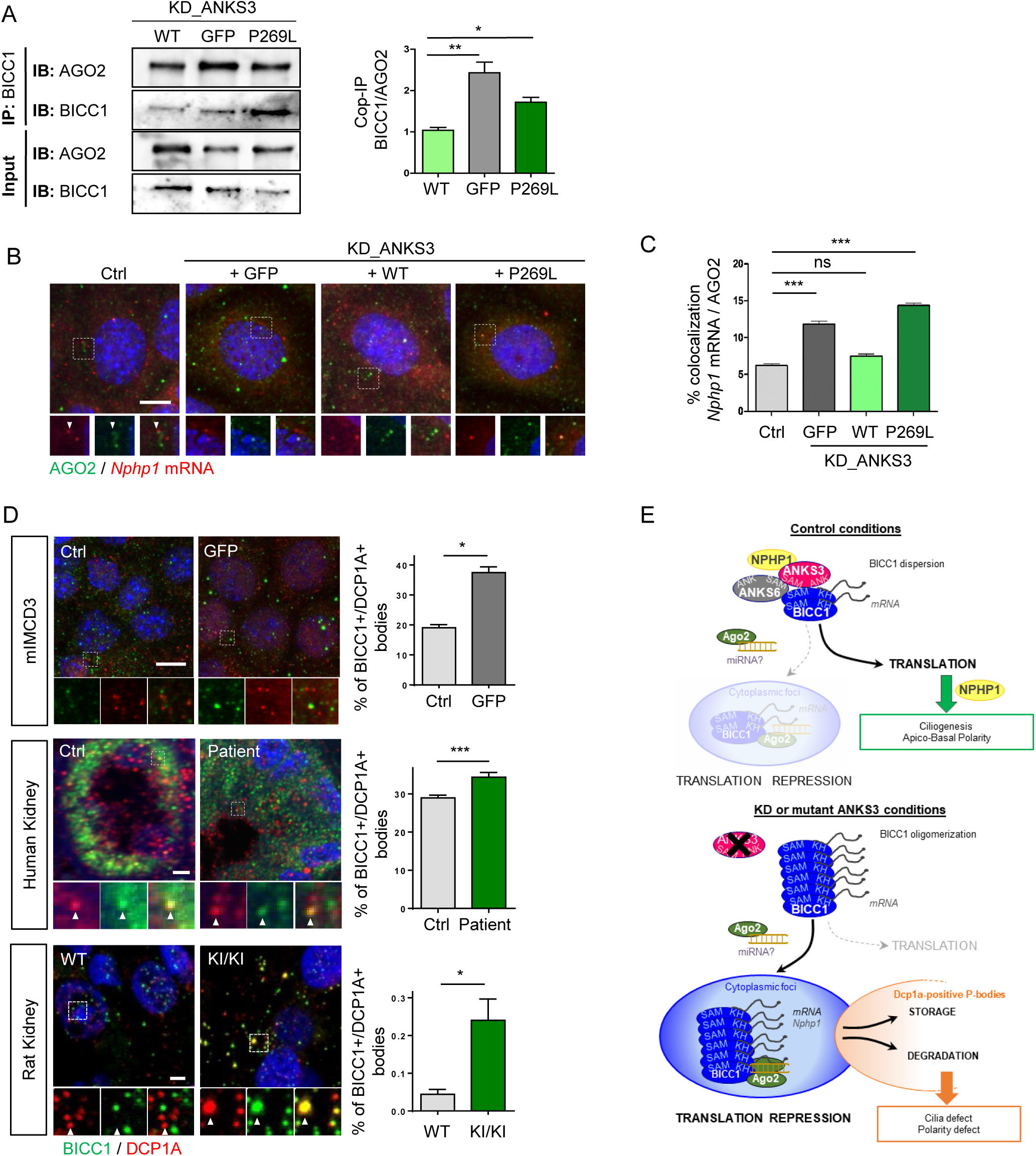
AGO2 interacts with BICC1 and colocalizes with *NPHP1* mRNA: **(A)** Lysates from KD_GFP, KD_WT, and KD_P269L mIMCD3 cells were immunoprecipitated with BICC1 antibody. Western blots of immunoprecipitates were performed using BICC1 and AGO2 antibodies and BICC1/AGO2 interaction was quantified. **(B-C)** Immunofluorescence of the *Nphp1* mRNA molecules with CY3-coupled smiFISH probes (red) and AGO2 protein (green) (B). Scale bar, 5µm. Quantification (%) of *Nphp1* smiFISH spots with AGO2 signal colocalisation is based on 3 independent experiments. Values are the mean ± sem. *p*-values were determined by one-way ANOVA followed by Dunn’s multiple comparison test; ***p≤0.001; ns: non-significant, p>0.05. **(D)** mIMCD3 (upper panel), human kidney biopsy (middle panel) and section of rat kidney (bottom panel) were stained for BICC1 (green) and the P-bodies marker DCP1A (red). Scale bar, 10µm. Quantifications of colocalization of BICC1 and DCP1A by colocalizer ImageJ protocol are performed with a distance of 3 px, and are based on 3 independent experiments. Values are the mean ± sem. *p*-values were determined by Mann-Whitney test; ***p≤0.001; *p≤0.05. **(E)** Schematic model of the proposed mechanism by which ANKS3-BICC1 complexes regulate post-transcriptional ciliary gene expression.

### BICC1 regulates *NPHP1* mRNA transcript stability by associating with AGO2

Previously, BICC1 was shown to modulate *Adcy6* and *Pkia* expression by controlling the mRNA translation together with the RISC complex. BICC1 promotes the loading of the DICER/miRNA precursor complexes onto its mRNA targets, which are then transferred to Argonaute 2 (AGO2) to be silenced (19). As AGO2 has also been implicated in miRNA-dependent mRNA degradation (34), we thus considered the role of AGO2 in the regulation of mRNA degradation by the ANKS3/BICC1 complex. We found that endogenous BICC1 and AGO2 proteins interacted (Figure 6A) in mIMCD3 cells, and that these interaction tend to be enhanced in both KD_GFP and KD_P269L cells (Figure 6A and Supplementary Figure 7A). These results suggested that ANKS3 could negatively control the BICC1/AGO2 interaction, subsequently inhibiting degradation of *Nphp1* transcript by the RISC complex. In favour of this hypothesis, we observed increased co-localization of *Nphp1* mRNA molecules with AGO2 in KD_GFP and KD_P269L cells (Figure 6B, C). We then investigated whether the downregulation of *Nphp1* transcript was mediated by AGO2. As expected, silencing of *Ago2* expression in mIMCD3 cells (Supplementary Figure 7B) resulted in an increased expression of *Nphp1* mRNA in both KD_GFP and KD_P269L cells (Supplementary Figure 7C). Finally, BICC1 was found to colocalize more frequently with DCP1a-positive processing bodies (where mRNA storage or degradation occurs) in KD_GFP cells, as well as in NPH patient kidney biopsy samples and rat kidney (Figure 6D).

Altogether, these data indicate that the impaired ANKS3-BICC1 interaction, due to the absence or variant of ANKS3, increases BICC1/AGO2 interaction and then the transfer of *Nphp1* transcript to the RISC complex for degradation in processing bodies.

## Discussion

Ciliopathies are a complex group of genetic diseases characterized by developmental or functional disorders of multiple organ systems. Although most ciliopathies result from pathogenic variants of structural ciliary proteins, in this study we decipher a novel mechanism involving a post-transcriptional regulator of ciliary gene expression, ANKS3, based on the identification of the P269L *ANKS3* variant in patients with nephronophthisis. This variant adversely affects cilium morphology and function, as well as epithelialization in kidney tubular cells *in vitro*. In addition, both zebrafish and rat models validate the pathogenic effect of this mutation and its impact on kidney function leading to urine concentration defect and tubular dilatation as observed in NPH. As a mechanism, we showed that the mutated ANKS3 fails to properly interact with BICC1, a known regulator of the translation of ciliary gene transcripts. By demonstrating (1) the key role of ANKS3 in regulating the stability of *Nphp1* transcript through modulation of BICC1 function and the recruitment of AGO2-RISC complex, (2) that the impairment of ANKS3/BICC1 complex activity has deleterious effects on the biogenesis and function of primary cilia, and (3) that *ANKS3* mutation is a genetic cause of NPH, we highlight ANKS3 as a novel renal ciliopathy gene and its involvement in a novel pathophysiological mechanism for NPH-related ciliopathies. Therefore, we propose *NPHP21* as an alias for *ANKS3*.

ANKS3 has been recently shown to negatively regulate BICC1 homo-polymerisation, and therefore the size of the BICC1 foci, *via* the formation of SAM domain-dependent ANKS3-BICC1 hetero-polymers (15). In agreement with this observation, we showed that in absence of ANKS3 or when ANKS3-BICC1 interaction is impaired, BICC1 forms larger foci in both mIMCD3 cells and patient kidney cells. We thus propose the following model (Figure 6E): alteration of ANKS3/BICC1 interaction leads to abnormal extension of BICC1 homo-polymers accompanied with enhanced ability to bind mRNA targets. This extended BICC1 homo-polymerisation leads to increased recruitment of AGO2-RISC complex, directing transcripts to DCP1a-positive P-bodies likely promoting their degradation. As a consequence, various cellular processes are impaired, notably epithelial morphogenesis and ciliary biogenesis and function. These results are in agreement with recent studies showing that BICC1 mediates decay of *Dand5* transcript in laterality organs (26, 27) and that ANKS3 negatively regulates BICC1 mRNA interaction (35, 36). Therefore our data highlight a new function of BICC1 in the regulation of mRNA stability in renal ciliopathies, as observed here for *Nphp1*. Also, it is not excluded that ANKS3 could directly acts on transcript regulation, as it has been recently shown that it binds RNA molecules, shared with BICC1, and associates to AGO2 (20). This could reveal a mechanism where *Nphp1* transcript bind both ANKS3 and BICC1 in a competitive complex that finely tune its stability by preventing BICC1 aggregation and subsequently its degradation through the RISC Ago2 complex.

As part of this novel mechanism of post-transcriptional regulation of NPHP1 and potentially other key ciliary transcripts, we propose that the ANKS3-BICC1-dependent mRNA regulation most probably occurs in the cytoplasmic foci in epithelial cells but also possibly in close proximity to the cilia, as it was previously suggested by the association of BICC1 with the centrosomal protein OFD1 (18). We show that the ciliary phenotypes observed in ANKS3 mutant conditions can be explained only in part by the dysregulation of the expression of NPHP1. This suggests that other ciliary (or non-ciliary) gene transcripts, that remained to be identified, could follow a similar ANKS3/BICC1 dependent regulation.

ANKS3 loss of function and p.P269L variant are responsible for opposite ciliary phenotypes (shorter *versus* longer cilia). A similar apparent discrepancy on cilia phenotypes has previously been observed with other ciliary proteins, including IFT components (IFT172, IFT54) or TZ proteins, for which loss-of function mutations led to shorter cilia whereas missense mutation lead to longer cilia (5, 11, 37). In contrast, loss of function of the TZ components NPHP1 and NPHP4 is associated with longer cilia in kidney tubular cells (this study, 12, 30).

Regarding the impact of ANKS3 on the composition of TZ, the fact that the amount of NPHP1 is decreased under both mutant and KD_ANKS3 conditions is consistent with the downregulation of the *Nphp1* transcript in mIMCD3 cells. Notably, the level of NPHP4 at TZ was not affected, which is expected because recruitment of NPHP4 to TZ does not require NPHP1 as shown in several cell types (38). In contrast, TMEM67 was absent from TZ in KD_ANKS3 cells while unchanged in P269L mutant cells. Since the recruitment of TMEM67 to TZ does not depend on the expression of NPHP1 (39), this suggests that loss of ANKS3 may also regulate broadly TZ component expression defect. Whatever the mechanism, loss of TMEM67 at TZ in KD_ANKS3 cells compared to mutant conditions, could explain the differential alteration of ciliary length and composition (39–41) and the fact that re-expression of NPHP1-V5 only restores the longer cilia in P269L mutant cells and not the shorter cilia in KD_ANKS3 cells.

In addition to TZ defects, it is quite remarkable that another ciliary phenotype observed in both KD and p.P269L ANKS3 conditions is the mislocalization of the InvsC components all along the axoneme. The mechanisms responsible for this relative elongation remain to be identified, ANKS3 is however known to interact with components of the InvsC including ANKS6 and NEK8 and may regulate their assembly and/or entry in the ciliary compartment. Notably, the InvsC appears to play a critical role in cilia-dependent signalling in the laterality organs as evidenced by the fact that laterality defects in association with kidney anomalies are recurrent phenotypes associated with variants of InvsC components genes in both murine models and humans (42–47). In agreement, *situs inversus* was associated with a previously described human *ANKS3* variant located in the Ankyrin repeats domain of the protein (28) as well as in *anks3* zebrafish models (this study and (14)). Laterality defects have not been reported in any of the siblings in the family described here, nor in our rat model. We can therefore assume that the p.P269L variant is a hypomorphic allele resulting mainly in a kidney-and liver-associated phenotype. In addition to the ANKS3-associated InvsC defects, *Bicc1* was also involved in laterality (22) which relies on its role on the regulation of *Dand5* mRNA decay in response to flow (26, 27). In this context, ANKS3-associated laterality defects could be attributed to its function in InvsC or/and BICC1-related functions.

Finally, in addition to ciliary defects, alteration of ANKS3 led to abnormal apico-basal polarity in epithelial cells. This result is in agreement with the role of ANKS3 in control of NPHP1 expression since loss of NPHP1 results in polarity defects (12). Interestingly, a previous study in *C. elegans* highlighted the function of ANKS3 and its binding partners (NEK8, NEK7, ANKS6 and INVS) in the control of actin cytoskeletal organization through the regulation of the RhoGTPase Cdc42 and its downstream effector ACK1 (48). Moreover, NPHP1 was previously shown to interact and colocalize with ACK1 at cell-cell junctions in kidney cells (49). We can thus hypothesize that disrupting the interactions between ANKS3 and its partners together with the decreased *Nphp1* expression, affect actin cytoskeletal organization and cell contractility, both of which are well-known modulatory factors of ciliogenesis and apico-basal cell polarity (50).

In conclusion, the data presented in this paper highlight the crucial functions of ANKS3 as a key regulator of BICC1 function in the control of the expression of a key cilia-related gene, orchestrating the biogenesis and function of primary cilia, and regulating intracellular signalling pathways required for epithelialization. Our findings provide evidence for a new mechanism for renal ciliopathies involving post-transcriptional dysregulation of ciliary gene expression, which may contribute significantly to kidney tubulo-interstitial lesions.

## Methods

### Study Approval

The human studies are kept in full accordance with the principles of the Declaration of Helsinki and Good Clinical Practice guidelines. The human renal biopsies and fibroblasts belonging to the Imagine Biocollection are declared to the French Minister of Research under the number DC-2020-3994 and this biocollection was approved by the French Ethics committee for research at Assistance Publique-Hôpitaux de Paris (CERAPHP) under the IRB registration number: #00011928. Animal studies: All animals were handled in strict accordance with good animal practice as defined by the national animal welfare bodies, and all animal procedures were performed in accordance with protocols approved by the ethical committee of the « Ministère de l’Enseignement Supérieur, de la Recherche et de l’Innovation » (N°APAFIS#22608-2018042715366694).

### Study participants

Genomic DNA was extracted from blood cells and fibroblasts were obtained from skin biopsies from patients using standard procedures. Kidney biopsies were evaluated by pathologists.

### Homozygosity mapping, exome sequencing, and mutation calling

Homozygosity mapping was performed using ‘Human Mapping 250k NspI’ array and parametric logarithm of odds scores were calculated with MERLIN software, assuming autosomal recessive inheritance.

For whole exome sequencing, genomic DNA isolated from blood lymphocytes was subjected to exome capture using Agilent SureSelect human exome capture arrays (Life Technologies), followed by next generation sequencing on the Illumina® sequencing platform. The raw data FASTQ files were first mapped to NCBI build 37/hg19 reference genome using Burrows-Wheeler Aligner (BWA). Downstream processing was carried out with Genome Analysis Toolkit (GATK), SAMtools, and Picard Tools, following documented best practices (http://www.broadinstitute.org/gatk/guide/topic?name=best-practices). All variants were annotated using a software system developed by the Paris Descartes University Bioinformatics platform. The mean depth of coverage obtained was greater than x72, and >94% of the exome was covered at least x15. Different filters were applied to exclude all variants located in non-exonic regions, pseudogenes, untranslated regions, or known polymorphic variants with a frequency above 1% annotated in databases such as dbSNP, 1000 genome projects, and gnomAD, including all variants identified by in-house exome sequencing (12,426 exomes and 1,355 ciliomes). The functional consequence of missense variants was predicted using SIFT (http://sift.jcvi.org/www/SIFT_enst_submit.html), PolyPhen2 (http://genetics.bwh.harvard.edu/pph2/) and CADD/PHRED2 (http://cadd.gs.washington.edu/) software. Traditional Sanger sequencing was performed using the primers Forward 5’-TCA GTT TGC CGT AAG GAT TAG T-3’ and Reverse 5’-CTA TTC CAT TCT TGC CAA CAC-3’ to validate the next generation sequencing findings and the segregation of the mutation in the family.

### Zebrafish experiments

Zebrafish were maintained at 28.5°C under standard protocols. The following transgenic lines were used: *Tg(bactin:arl13b-GFP)^hsc5Tg^*, (51) *Tg(sox17:GAL4;cmlc2:GFP)* (gift from M. Bagnat., Duke University, USA) and *Tg(UAS:GCaMP6f; cryaa:mCherry)^icm06^*(52). To generate the *anks3* mutant fish line, TALENs targeting exon 3 were designed and cloned into pCS2TAL3RR or pCS2TAL3DD expression vectors using Golden Gate assembly (53, 54). RNA encoding each TALEN arms (left arm target, 5’-CTG ATG TAC GCT GCT T-3’; right arm target 5’-ACA TCG CTA ACC TGC TG CT-3’) were synthesized using the mMESSAGE mMACHINE kit (Ambion) and injected into one-cell stage embryos at 100 pg per embryo. Mutant alleles were identified by high-resolution melt analysis of PCR products generated with the following primers: forward, 5’-GGA GAT GTG GAT TTG GAT GG-3’ and reverse, 5’-CTG CAA GAG GAA GTG AGC AA-3’. Genotyping of the *anks3^ii002^* allele was performed using allele-specific primers: forward 5’-TAC GCT GCT TAC ATC GGT CA-3’ for the WT allele, forward 5’-ACG CTG CTT ACA TAT CTG CTG-3’ for the mutant allele, and reverse, 5’-TGC ACA GAT CAA TTG TTC ACC-3’. For RNA rescue experiments, WT and mutated human ANKS3-GFP mRNA were synthesized using the mMESSAGE mMACHINE kit and injected at 150 pg into one-cell stage embryos. Heart looping phenotypes were analysed using a Leica M165FC stereoscope. For live imaging of *Tg(bactin:arl13b-GFP)* or *Tg(sox17:GAL4;cmlc2:GFP); Tg(UAS:GCaMP6f; cryaa:mCherry)* transgenic animals, all acquisitions were performed using a Zeiss LSM confocal microscope after immobilization of live embryos in 0.8% low melt agarose in glass-bottom Petri dishes (Ibidi). Z-stacks were performed to acquire the whole Kupffer’s vesicle and projections were assembled and analysed using ImageJ software to determine cilia range of motion (as described previously (51)), and GCaMP signal intensity.

### Production of Anks3 knock-in rats

To generate the humanized p.P269L Anks3 knock-in rat strain, the complex sgRNA /Cas9protein/ and the donor oligonucleotide sequence was microinjected into the pronuclei of rat zygotes of the Sprague-Dawley strain. The donor oligonucleotide sequence (gCCACGAGGGGcctCGCGCCC) carried the mutation P269L and additional silencing mutations to prevent the cutting of the modified sequence by the guide sequence DNA oligonucleotides. Zygotes surviving microinjection (∼75%) were implanted into pseudopregnant females. Newborns were analyzed by PCR amplifying (∼300bp) the targeted sequence (Forward: 5’-gctgtaagttcaaggcctgcct-3’, Reverse: 5’-gagccagacggggactaaaacc-3’), followed by DNA denaturation, renaturation and incubation with T7 exonuclease that will cleave DNA hybrids containing mismatches. Founders with insertion donor sequences were selected to generate lines of homozygous rats after 4 backcross to Sprague-Dawley strain for phenotyping. DNA sequences of WT and *Anks3* mutant (KI) rats and corresponding amino acids: WT - ATC CAC GAG GGG CCT CGC GCC CTG GCC AGG ATC , KI - ATC CAC GAG GGG C**TG** CG**G** GCC CT**C** GCC AGG ATC, corresponding to amino acid sequence I H E G P/**L** R A L A R I.

### Histology and analysis of kidney lesions

Kidneys were fixed in formalin and paraffin embedded. Three-micrometer–thick sections were stained with hematoxylin-eosin (HE) and picrosirius red. Interstitial fibrosis was assessed on sirius red-stained paraffin sections (20 X) under polarized light on 8 different cystic WT and KI/KI scanned whole kidneys. It was quantified using computer-based morphometric analysis software (Axioplan, Axiovision, Zeiss, Germany). Data are expressed as percentage of positive areas examined. Determination of cyst percentage was performed on HE-stained whole kidney sections. Microphotographs from 8 WT and 8 KI/KI kidneys were analyzed with NIH ImageJ software.

### RNA extraction and RT-qPCR

Total mRNA was isolated using Qiagen Extraction Kit and then treated with DNaseI. 1.5 µg of total RNA was reverse-transcribed using Superscript II (Life Technologies). Relative expression levels of genes of interest were determined by real-time RT-PCR using the Absolute SYBR Green ROX Mix (ABgene) and specific primers (Table S1). Experiments were repeated at least 3 times and gene expression levels were normalized to *GAPDH* or *HPRT*. Data were analysed using the 2^−ΔΔCt^ method (11).

### Human primary fibroblasts and mIMCD3 cell culture

Control and affected patient fibroblasts were cultured in Opti-MEM supplemented with 10% foetal bovine serum, 100 U ml−1 penicillin and 100 mg ml−1 streptomycin (all from Life technologies). Control and NPH patient fibroblasts (1.5×10^4^ or 2.5×10^4^ cells, respectively) were plated on coverslips and grown for 2 days (low confluence) or 6 days followed by 48 h serum deprivation (high confluence) before fixation. Murine inner medullary collecting duct (mIMCD3) cells were cultured in DMEM GlutaMaxI medium containing 10% fetal bovine serum (FBS), 100 U ml−1 penicillin and 100 mg ml−1 streptomycin (all from Life technologies). For immunofluorescence, 2.5×10^4^ cells were plated on coverslips and grown for 4 days before fixation. For all experiments, the level of confluence was checked visually and cells were counted using Beckman Coulter instrument to ensure similarity between control and samples. All cells were regularly tested for mycoplasma contamination and were mycoplasma-free.

### Plasmids and lentiviral infections

Anks3-knockdown was performed in mIMCD3 cells by lentiviral infection with a shRNA expressing construct cloned in Lv122-puromycin vector (Gene Copoeia EX-H2236-Lv122). Stable cell lines re-expressing WT or mutant ANKS3-GFP forms were obtained using lentiviral infection of an ANKS3-GFP construct cloned in Lv122-puromycin vector. ANKS3-GFP variants were obtained through site-directed mutagenesis using a Pfu turbo kit (Invitrogen). NPHP1-V5-rescued cell lines were obtained by lentiviral infection with a NPHP1-6A construct tagged with V5 (55) cloned in the pRRL.SIN.cPPT.PGK/GFP.WPRE vector (Adgene).

### Short interfering RNA (siRNA) transfection

SiRNA transfection were performed on cells in suspension using the Lipofectamine 2000 transfection kit according to the manufacturer’s instructions. Cells were transfected with 4 μl of 2 µM of siRNAs listed in Table S1. Cells were collected 72 h after transfection for analysis.

### Antibodies

The following antibodies were used in the study: acetylated α-tubulin (Sigma, 6–11-B-1, 1:10,000), acetylated α-tubulin (Abcam, ab24610, 1:500), α-tubulin (Abcam, ab18251, 1:500), ARL13B (Proteintech, 17711-1-AP, 1:400), GP135 (R&D, AF1556, 1:200), Polycystin-2 (Santa Cruz, SC10376, 1:50), NPHP1-C20 (Santa Cruz, 1:200), NPHP1 (BiCell, 90001, 1:150), Rabbit NPHP1 antibody was described previously (55) and used at 1:500 dilution for western blots, NPHP4 (BiCell, 90004, 1:150), TMEM67 (BiCell, 90103, 1:200), DCP1A (Abnova, 55802, 1:200), γ-tubulin (Millipore, C-20, 1:200), γ-tubulin (Sigma, DQ-19, 1:500), γ-tubulin (Sigma, GTU-88, T6557, 1:500), ANKS3 (Sigma, HPA041409, 1:50 for IF and 1:1000 for WB), ANKS6 (Sigma, HPA008355, 1:200), BICC1 (Sigma, HPA045212, 1:50 for IF and 1:1,500 for WB), GAPDH (Millipore, MAB374, 1:4,000), DsRed (Clontech, 632496, 1:4000), NEK8 (gift from D. Beier 1:500), chicken GFP (2B Scientific, 1020, 1:200), Smo (Abcam ab38686 1:100), AGO2 (Sigma SAB4200085 1:100 for IF and 1:1,000 for WB), ZO-1 (Sigma, R40.76, MABT11, 1:100), Myc (Thermo Fisher MSP139P1 1:5,000 for WB), and V5 (AbD Serotec MCA1360 1:5,000). Highly cross-adsorbed secondary antibodies (Alexa Fluor 488, Alexa Fluor 546, AlexaFluor 555, AlexaFluor 532, and Alexa Fluor 647) were obtained from Molecular Probes (Life Technologies) and used at 1:400 dilution.

### Immunofluorescence staining

Fibroblasts and mIMCD3 cells were fixed in 4% PFA in 1X PBS followed by treatment with 50 mM NH_4_Cl and permeabilization with 0.2% Triton-X 100 or fixed in ice-cold methanol for 5 min. Cells were then blocked with 1% skim milk or 3% BSA containing 0.1% Triton-X100 before incubation with primary antibody (1-3 h at room temperature or overnight at 4°C) and Alexa Fluor-conjugated secondary antibody (30 min at room temperature). Appropriate controls were performed omitting the primary antibodies. DNA was stained with DAPI or Hoechst (Life Technologies). Confocal images were obtained on LEICA SP8 microscope with a x40 (HC PL APO 40x/1.30 Oil CS2) or x63 (HC PL APO 63x/1.40 Oil CS2) objectives and Leica LAS-AF software. Images were analysed with ImageJ software. Z-stacks were acquired with identical acquisition settings (gain, offset, laser power) and all measurements of fluorescence intensity were performed on sum intensity projection.

Formalin-fixed, paraffin-embedded human tissue biopsies were sectioned (8 μm thickness) using a Leica microtome. Sections were immersed in xylene baths (5 min each) then rehydrated for 5 min in ethanol baths of decreasing concentration (100%, 95%, 70%, and 40%) and finally immersed in MilliQ water for 5 min. Dako target retrieval solution (Dako ref. S1699) was used according to the manufacturer’s instructions. Slides were blocked for 45 min at 4°C by 10% normal donkey serum diluted in PBT (DPBS with 0.1% Triton X100). Primary and secondary antibodies were used as described above. Slides were mounted in adapted medium and analysed under an inverted confocal microscope (Zeiss LSM 700).

Rat kidneys were fixed in formalin and paraffin embedded. Three-micrometer–thick sections were deparaffinized in 2 BioClear baths (10 min each) then rehydrated for 5 min in ethanol baths of decreasing concentration (100%, 90%, 80%) and finally immersed in MilliQ water for 5 min. Dako target retrieval solution (Dako ref. S1699) was used according to the manufacturer’s instructions. Slides were blocked for 20 min at RT in TBST-BSA 3% and FBS 10%. Primary and secondary antibodies were used as described above. Slides were mounted in Mowiol medium and analysed under an inverted confocal microscope (Leica SP8)

### Electron microscopy

For scanning electron microscopy, cells were washed in PBS, fixed with 2.5% glutaraldehyde, and processed as reported previously (56). Cells were visualized using a scanning electron microscope (Jeol. 6700F).

### Measurement of cell height

Cell height was measured on mIMCD3 cells grown on filter stained with β-catenin that label cell membrane. The height was evaluated on orthogonal views on ImageJ software. Quantification of the cell height was done as the distance from the base to the top of the cells.

### Measurement of the length of cilia and the Inversin compartment by IF

Cilia length was measured on slides stained with acetylated-tubulin antibody. The length was evaluated on maximum projection on ImageJ software. The measurement of the size of the inversin compartment was evaluated as the ratio of the length of INV, NEK8 or ANKS6 staining and the total cilia length (acetylated αtubulin staining) on maximum projection on ImageJ software.

### Quantification of transition zone proteins fluorescence intensity

The mean intensity of NPHP1, TMEM67 and NPHP4 fluorescence was measured on sum projection of 14 z-stacks. Identically sized concentric rings encompassing the transition zone were used as the area of reference for all transition zone fluorescence intensity measurements. Image quantification was performed using ImageJ.

### Colocalization analysis

Post-acquisition colocalization image analysis was performed using Icy software (57). The undecimated wavelet transforms Spot Detector plugin and the Colocalizer protocol were used to quantify the number of particles and colocalized particles, respectively.

### Protein extraction and western blot

Cells were lysed in 50 mM Tris-HCl, 150 mM NaCl, 0.5% sodium deoxycholate, 2 mM EDTA, 1% Triton X-100, 0.1% SDS, and protein concentrations were determined using a BCA protein assay kit (ThermoScientific). 30-50 µg of protein were loaded on 8% or 10% acrylamide gels, blotted on PVDF membranes (Millipore). Membranes were blocked in 5% milk for one hour at room temperature and incubated and incubated using the indicated antibodies. Western blots were then analysed with Bioprofil software.

### Coimmunoprecipitation

*Miltenyi beads:* ANKS3-null mIMCD3 cells were transiently transfected using the calcium phosphate method with different tagged protein constructs. After 48 h, cells were harvested in accordance with the Miltenyi Biotec beads protocol. Protein concentrations were determined using a BCA protein assay kit (ThermoScientific).

*Sepharose beads:* 48 h post-transfection, cells were lysed in 50 mM Tris-HCl pH 7.5, 150 mM NaCl, 0.5% Triton and lysates were incubated with rabbit or mouse isotypic control antibodies and G-protein beads (Sigma) for 1 hr at 4°C. Pre-cleared lysates (containing 1 mg of protein) were then incubated with mouse monoclonal anti-GFP antibodies (Roche) coupled to G-protein beads for 3 h at 4°C. Beads were then washed three times with increasing amounts of NaCl (500 nM; 300 nM and 150 nM in 50 mM Tris-HCl pH 7.5), resuspended in 2X sample buffer (Sigma), and boiled for 5 min.

### Calcium switch assay and TER measurement

mIMCD3 cells grown on 65 mm Transwell filters (Corning) for 7 days were subjected to Ca^2+^ switch as described by (58). Briefly, cells were washed with PBS containing 4 mM EGTA and calcium-free DMEM (Life Technologies) containing 4 mM EGTA was then added to the cells for 45 min to disrupt cell junctions. Cells were then washed twice with normal culture medium (DMEM/F12). TER was determined using a Millicell-ERS volt–ohm meter (Millipore) immediately after addition of normal growth medium and at the indicated time points. 6 h after Ca^2+^ switch, cells were fixed with 4% PFA and processed as described above.

### mRNA stability assays

mIMCD3 cells were treated with 1 μM actinomycin D (Sigma-Aldrich) to block mRNA synthesis. Cells were harvested at various time points after actinomycin D treatment. *Nphp1* and *Gapdh* mRNA levels were quantified by quantitative RT-PCR and mRNA half-lives were calculated from plots of mRNA levels as a function of time. We also performed the experiments in the presence of DMSO as a control.

### RNA-immunoprecipitation (RIP) assay

RIP assays were performed on cell extracts from mIMCD3 control cells, mIMCD3 shANKS3 cells re-expressing GFP plasmid, mIMCD3 shANKS3 cells re-expressing WT ANKS3, and mIMCD3 shANKS3 re-expressing P269L mutant ANKS3 with anti-BICC1 polyclonal antibody using the RiboCluster Profiler RIP-Assay Kit (MBL, Japan, RN1001) according to manufacturer’s recommendations. Briefly, RNA-protein complexes were pulled-down with 5 µg of rabbit anti-BICC1 antibody. Then, bound RNAs were recovered from RNA-protein complexes, used as templates for cDNA synthesis, and analysed by classical PCR followed by agarose gel electrophoresis. Relative expression levels of genes of interest were determined by semi-quantitative RT-PCR (25 cycles for ADCY6 and GAPDH and 30 cycles for NPHP1, primers are listed in Table S1) using the Green Master Mix (Promega) specific primers.

### SmiFISH (59)

Primary smiFISH probes and FLAPs (secondary probes, fluorescent conjugated) were produced and purchased from Integrated DNA Technologies (IDT). Primary probes were generated by high-throughput oligonucleotide synthesis in 96-well plates. To make use of low-scale synthesis (25 nmol), the total length of primary probes (transcript-binding + FLAP-binding) should not exceed 60 nucleotides. The secondary probes were conjugated to Cy3 or Cy5 via conventional 5′ and 3′ amino modifications. mIMCD3 cells were fixed in 4% PFA in 1X PBS followed by incubation in ice-cold at -20°C methanol overnight. Cells were then incubated with primary antibody and smiFISH probes in hybridization buffer (overnight at 37°C). Alexa Fluor-conjugated secondary antibody were added (30 min at room temperature) and DNA was stained with DAPI or Hoechst (Life Technologies). Confocal images were obtained on LEICA SP8 microscope. Images were analysed with ImageJ.

### Statistical analyses

Data are presented as the mean +/- sem. Means of two conditions were compared with Mann-Whitney test. Means of more than two conditions were analysed by one-way ANOVA followed by Dunn’s multiple comparison tests (GraphPad Prism V6). All results were obtained in at least 3 independent experiments. *p*<0.05 was considered statistically significant. No samples were excluded from the analysis. All image analyses and zebrafish and rat phenotypic analysis were performed in a blinded fashion.

### Availability of data

The data that support the findings of this study are available from the corresponding author upon reasonable request.

## Author Contributions

A.M and G.O. succeeded each other over the period of this study, they performed cell biology and biochemical experiments with the help of R.R., V.G, M.M and F.S. G.H. and L.D performed zebrafish studies under the supervision of M.D. A.M., A.V., B.L, S.H and R.P performed rat studies. B.K. recruited subjects and/or provided clinical information. C.H., E.F., P.K., A.W.S, P.S-M. and S.S. performed linkage and mutational analysis. SEM microscopy was performed by G.O. and T.B., with the input of P.B. G.O performed smFISH experiments under supervision of M. D and F.M. A.B., M. D. and S.S. designed the experiments and supervised the analyses. G.O., A.M., C.J., A.B., M.D. and S.S. wrote the manuscript (with input from co-authors).

## Acknowledgements

We are grateful to the patients for their participation. We thank C. Wyart, A. Prendergast and M. Bagnat for providing us with the *UAS:GCaMP6f* and *sox17:GAL4* zebrafish transgenic lines, respectively. We thank D. Stainier for supporting the generation of *anks3* TALEN line. We thank B. Schermer for providing us with the NPHP1 antibodies. We thank I. Anegon, S. Menoret and L. Tesson (Platform TRIP, Nantes) and J.P. Concordet (Platfrom Tacgene, Muséum d’histoire naturelle, Paris) for generation of the *Anks3* knock-in rat model. We thank A. Becker-Heck and A. Bizet for technical support. We greatly acknowledge N. Goudin and M. Garfa-Traoré (Necker cell imaging facility) for providing expert knowledge on confocal microscopy, E. Oakley, B. Linghu and F. Yang (NVS sequencing team), C. Bole-Feysot and M. Parisot (Imagine Genomic plateform) and the bioinformatic Plateform (Université Paris Cité, Institut Imagine). We thank the histology facility (S.F.R. Necker INSERM US24, Paris, France). We thank S. Berissi (Histology facility), the pathology department of Necker hospital, O. Pellé (Cell sorting facility), for their excellent technical assistance and LEAT (Imagine Institut). The authors are grateful to the following scientists for their support: Marie-Claire Gubler for her precious help with interpretation of kidney biopsies, H. Le Hir for providing intellectual input at various stages of the project. Finally, the authors thank C. Antignac for her precious help in manuscript correction.

---

**Supplementary Fig. 1.**
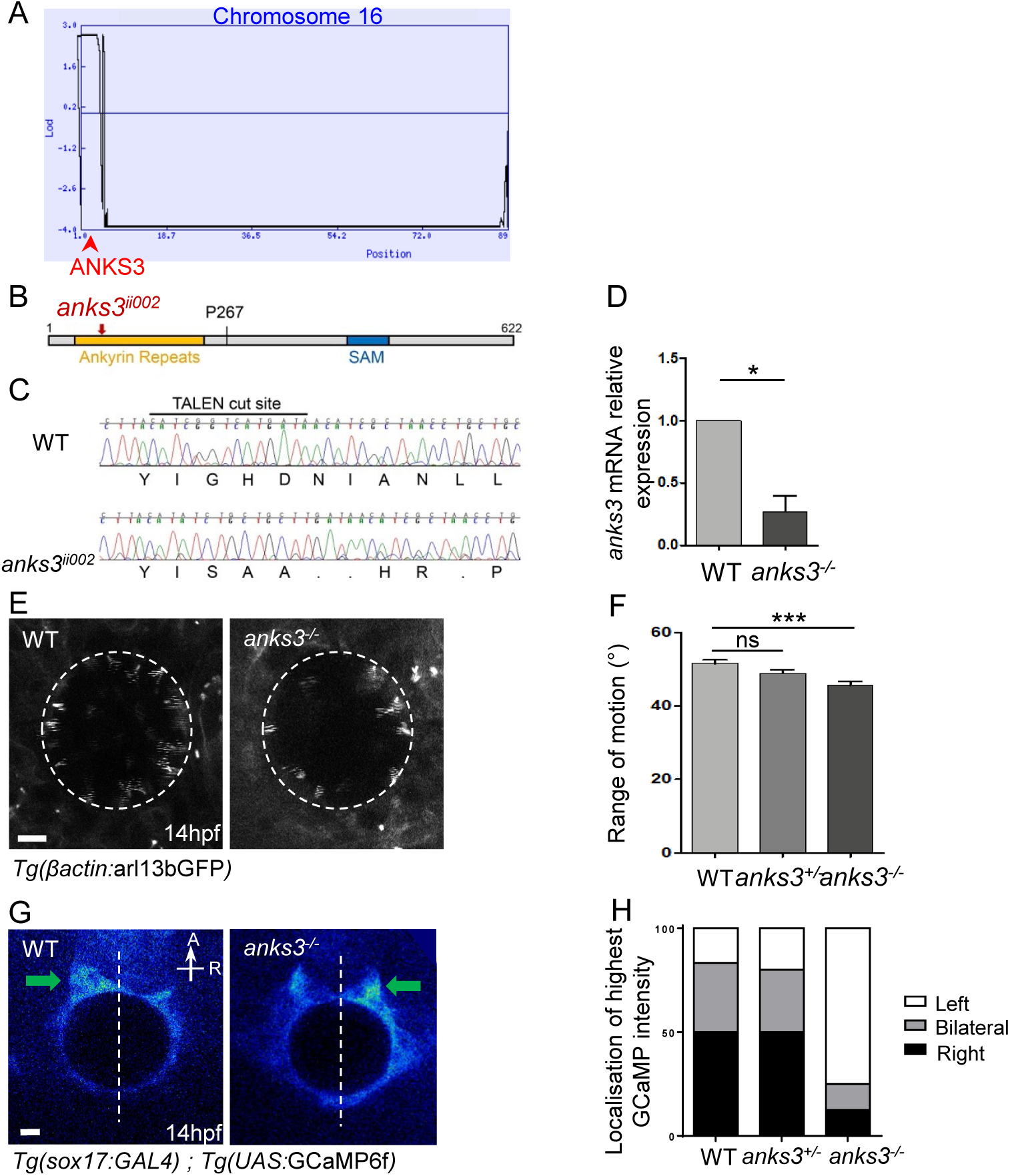
*ANKS3* mutation in three NPH and Zebrafish *anks3* mutant embryos exhibit abnormal cilia motility and Ca^2+^ signalling in Kupffer’s vesicle. **(A)** Linkage analysis revealed a homozygous region of 5.9 Mb on chromosome 16 from genomic position 16:113,599 (rs11248850) to 16:6,035,402 (rs2059271). A maximum logarithm of odds (LOD) score of 2.6577 was obtained at the *ANKS3* locus (red arrowhead). **(B)** Schema of the Anks3 protein showing the position of the TALEN-induced mutation, *anks3^ii002^*. **(C)** Chromatograms of the TALEN target region showing DNA sequences of WT and *anks3^ii002^* mutant alleles and corresponding amino acids. *anks3^ii002^* is an indel mutation (c.C246-251delCGGTCAinsATCTGCTGCT) that results in a frameshift and premature codon stop (p.Gly83Serfs86*). **(D)** Quantification of *anks3* mRNA relative expression by qPCR in WT and *anks3-*/- embryos at 48hpf. Values are mean ± sem.; *p<0.05 by t-test. **(E-F)** Live confocal imaging of Kupffer’s vesicle motile cilia in *Tg(bactin:arl13bGFP)* transgenic WT and *anks3* mutant embryos at 14hpf (E). Slow acquisition speed of motile cilia produces imaging artefacts, delineating the range of motion of each cilium. Quantification of the angle defined by the range of motion of motile cilia in WT (n=17), heterozygous (n=11), and mutant (n=10) embryos at 14hpf (F), as described in (E). Values are mean ± sem.; ***p<0.001, ns p>0,05 by Kruskal-Wallis’ test. Scale bars, 10µm. **(G-H)** Live imaging of cytoplasmic Ca^2+^ influx in WT and *anks3* mutant embryos at 14 hpf (G) using the calcium indicator GCaMP6f specifically expressed in Kupffer’s vesicle lining cells (sox17 promoter). Dorsal views, anterior (A) at top. Green arrows show the Ca^2+^ influx. Quantification (%) of the localization of the highest GCaMP6f intensity in WT (n=6), heterozygous (n=10), and *anks3* mutant (n=8) embryos at 14 hpf (H). Scale bars, 10µm.

**Supplementary Fig. 2.**
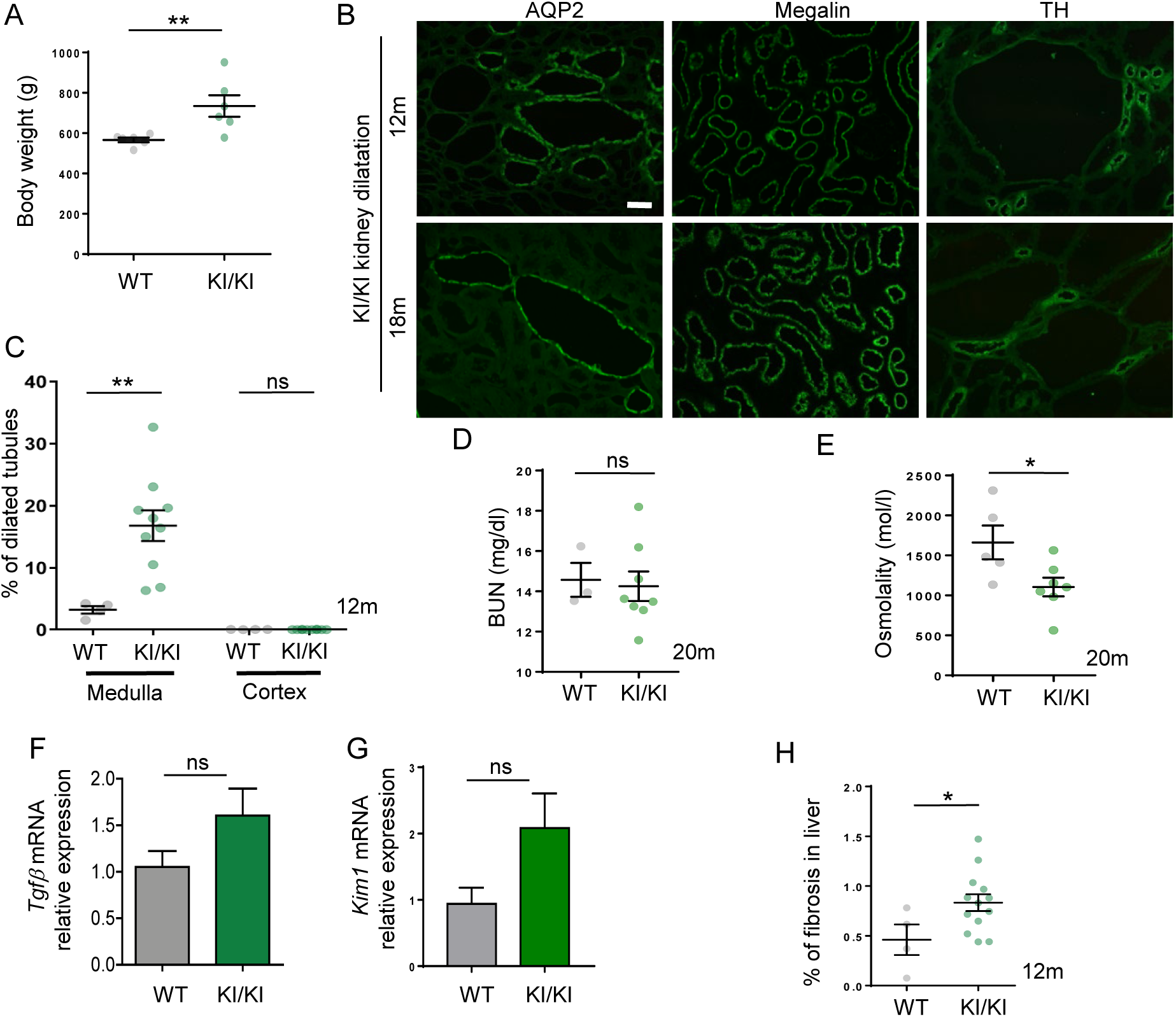
Analysis of the KI rat kidney. **(A)** Quantification of the body weight of 18-20 month old rats KI (n=5) and WT (n=5). Error bars represent the sem. p-values were determined with Mann-Whitney test. **p≤0.01 **(B)** Immunostaining of cross section of mutant rat kidney at 12 and 18 months for AQP2, TH (Tamm-horsfall) and megalin. Scale bar 20µm. **(C)** Quantification of percentage of dilated tubules in the medulla and the cortex of rat kidney at 12 months for WT (n=4) and KI/KI (n=10). Error bars represent the sem. p-values were determined with Mann-Whitney test. **p≤0.01; ns, not significant p>0.05. **(D, E)** Quantification of plasma blood urea nitrogen (BUN) (D) and urine osmolality (E) of WT and KI/KI rats at 20 months. Values are the mean ± sem. of n>3 animals. *p*-values were determined with Mann-Whitney test; **p≤0.01; *p<0.05; ns, not significant *p*>0.05. **(F, G)** qPCR analyses of *Tgfß* (F) and *Kim1* (G) mRNA relative expression in rat kidney at 18-20 months. Values are the mean ± sem. from 3 animals. *p*-values were determined by Mann-Whitney test. ns, not significant p>0.05 **(H)** Quantification of the percentage of fibrosis in rat liver at 12 months for WT (n=4) and KI/KI (n=13). Error bars represent the sem. p-values were determined with Mann-Whitney test. *p<0.05.

**Supplementary Fig. 3.**
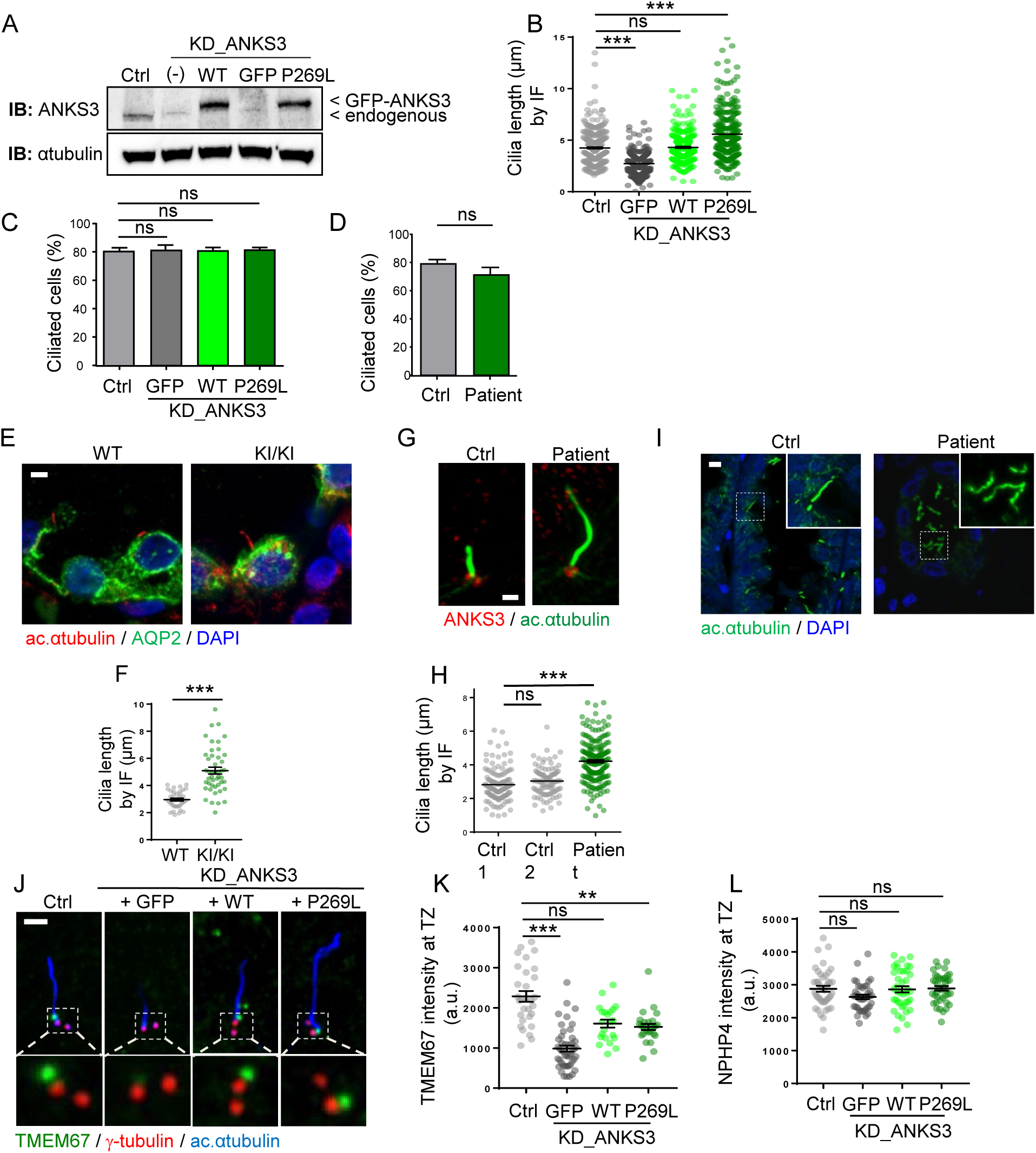
NPH-associated p.P269L mutation does not affect the percentage of ciliated cells but alters cilium length. **(A)** Immunoblot analysis of ANKS3 expression in control, KD_ANKS3, WT, GFP and P269L mIMCD3 cells (α-tubulin was used as protein loading control). **(B)** Quantification of the ciliary length by IF from Figure 2F. Values are the mean ± sem. of n>100 cells from 3 independent experiments; *p*-values were determined by one-way ANOVA followed by Dunn’s multiple comparison test; \*\*\**p*≤0.001; ns, not significant *p*>0.05. **(C-D)** Quantification of ciliogenesis (% of ciliated cells) in control, KD_ANKS3, WT, and P269L mIMCD3 cells (C) and NPH patient fibroblasts (D). Values are the mean ± sem. of 3 independent experiments with n >100 cells; *p*-values were determined by one-way ANOVA followed by Dunn’s multiple comparison test for (C) and Mann-Whitney test for (D); ns, not significant *p*>0.05. **(E)** Cilia staining (acetylated α-tubulin , red) in collecting duct (AQP2, green) in cross sections of rat kidney WT or KI/KI at 18 months. Scale bar, 3 µm. **(F)** Ciliary length in collecting duct in cross section of rat kidney at 18months Values are the mean ± sem. *p*-values were determined by one-way ANOVA followed by Dunn’s multiple comparison test; ***p≤0.001. **(G)** Immunofluorescence of ANKS3 (red) and acetylated α-tubulin (green) in control and patient fibroblasts. Scale bar 1 µm. **(H)** Quantification of the ciliary length by IF in NPH patient fibroblasts. Values are the mean ± sem. of n>100 cells from 3 independent experiments; *p*-values were determined by one-way ANOVA followed by Dunn’s multiple comparison test; \*\*\**p*≤0.001; ns, not significant *p*>0.05. **(I)** Cilia in kidney biopsies stained for acetylated α-tubulin (green). Scale bar, 5 µm. **(J)** Immunostaining of TMEM67 (green), γ-tubulin (red) and acetylated α-tubulin (blue) in control, KD_GFP, KD_WT, and KD_P269L mIMCD3 cells. Scale bar, 1µm**. (K-L)** Quantification of TMEM67 (K) and NPHP4 (L) fluorescence intensity at the TZ is based in 3 independent experiments. Values are the mean ± sem. of n>50 cells from 3 independent experiments; *p*-values were determined by one-way ANOVA followed by Dunn’s multiple comparison test; \*\*\**p*≤0.001; \*\**p*≤0.01 ns, not significant *p*>0.05.

**Supplementary Fig. 4.**
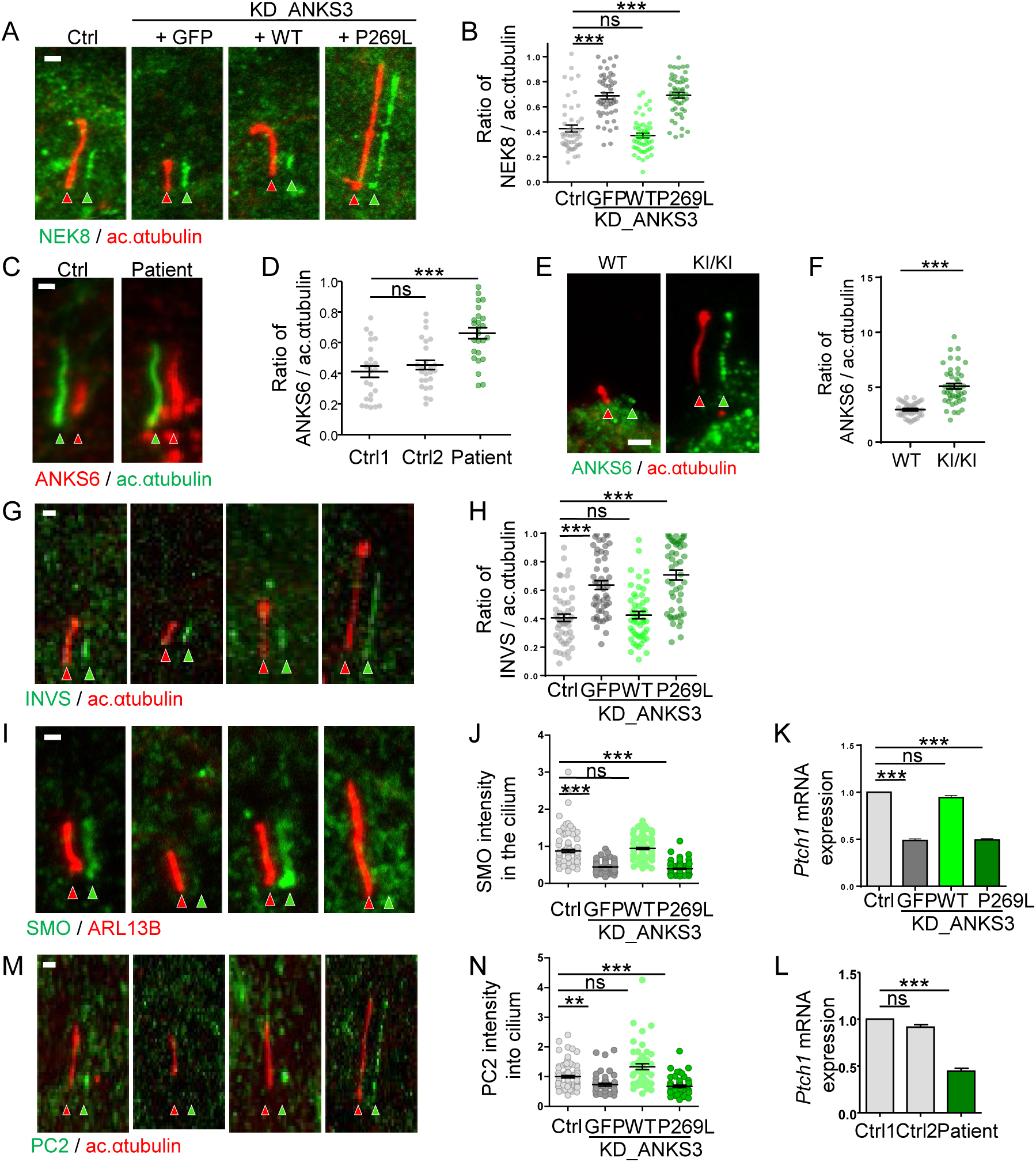
Analysis of ciliary proteins and signalling pathways in mIMCD3 cells and patient fibroblasts. **(A-B)** Ciliary distribution of NEK8 in control and KD_GFP, KD_WT, and KD_P269L mIMCD3 cells. Quantification of the size of the InvsC in mIMCD3 cells (B). Values are the mean ± sem. of n>100 cells from 3 independent experiments; *p*-values were determined by one-way ANOVA followed by Dunn’s multiple comparison test; \*\*\**p*≤0.001; ns, not significant *p*>0.05. **(C-D)** Ciliary distribution of ANKS6 in patient fibroblasts (C). Quantification of the size of the InvsC in NPH patient fibroblast (D). Values are the mean ± sem. of n>100 cells from 3 independent experiments; *p*-values were determined by one-way ANOVA followed by Dunn’s multiple comparison test; \*\*\**p*≤0.001; ns, not significant *p*>0.05. **(E-F)** Ciliary distribution of ANKS6 in collecting duct in cross section of rat kidney at 18months, stained for ANKS6 (green) and acetylated α-tubulin (red) (E). Scale bar, 1µm. Quantification of the size of the InvsC (F) in KI/KI and WT rats.**(G-H)** Ciliary distribution of INVS in control, KD_GFP, KD_WT, and KD_P269L mIMCD3 cells (G). Quantification of the size of the InvsC in mIMCD3 cells (H). Mean ± sem. of n>100 cells from 3 independent experiments, ns: non-significant, ***p<0.001, by two-way ANOVA and Dunn’s *post-hoc* test. **(I-J)** Ciliary distribution of smoothened (SMO, green) in cilia (ARL13B, red) in control, KD_GFP, KD_WT, and KD_P269L mIMCD3 cells treated with Smoothened Agonist (SAG) (I). Quantification of SMO fluorescence intensity in cilia (J). Values are the mean ± sem. of n>100 cells from 3 independent experiments; *p*-values were determined by one-way ANOVA followed by Dunn’s multiple comparison test; \*\*\**p*≤0.001; ns, not significant *p*>0.05. **(K-L)** qPCR analyses of relative *Patched1* mRNA expression in SAG-treated mIMCD3 cells (K) and patient fibroblasts (L). Values are the mean ± sem. from 3 independent experiments. *p*-values were determined by two-way ANOVA followed by Dunn’s multiple comparison test. ns: non-significant, ***p≤0.001. **(M-N)** Ciliary distribution of PC2 in mIMCD3 cells stained for PC2 (green) and acetylated α-tubulin (red) (M). PC2 staining intensity was quantified inside the ciliary region of interest (N). Mean ± sem. of n>100 cells from 3 independent experiments, ns: non-significant; **p<0.01, ***p<0.001, by two-way ANOVA and Dunn’s *post-hoc* test. Scale bars, 1µm.

**Supplementary Fig. 5.**
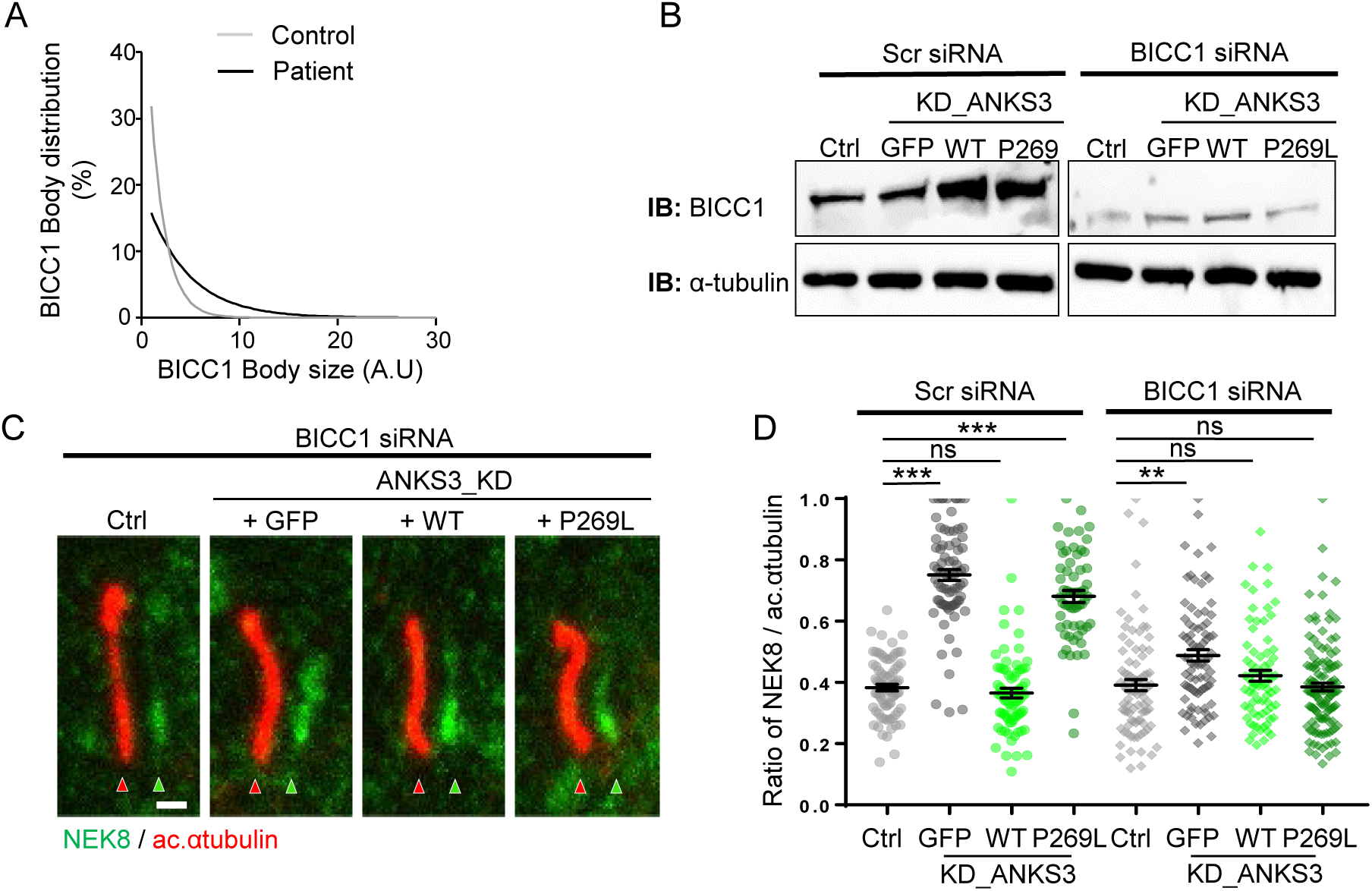
BICC1 knockdown rescues KD_ANKS3 and mutant phenotypes. **(A)** Distribution of size of BICC1 bodies using ImageJ spot detector protocol in NPH patient fibroblasts. **(B)** Western blot showing the efficiency of the BICC1-targeting siRNA in control, KD_ANKS3, WT, and mutant mIMCD3 cells (α-tubulin was used as protein loading control). **(C)** Ciliary distribution of NEK8 (green) in control, KD_ANKS3, WT, and P269L mIMCD3 cells transfected with scramble or BICC1-targeting siRNA. Scale bar, 1 µm. **(D)** Distribution of NEK8 was quantified as the ratio of NEK8 staining length to overall cilia length. Mean ± sem. from 3 independent experiments; ns: non-significant; **p<0.01, ***p<0.001, by two-way ANOVA and Dunn’s *post-hoc* test.

**Supplementary Fig. 6.**
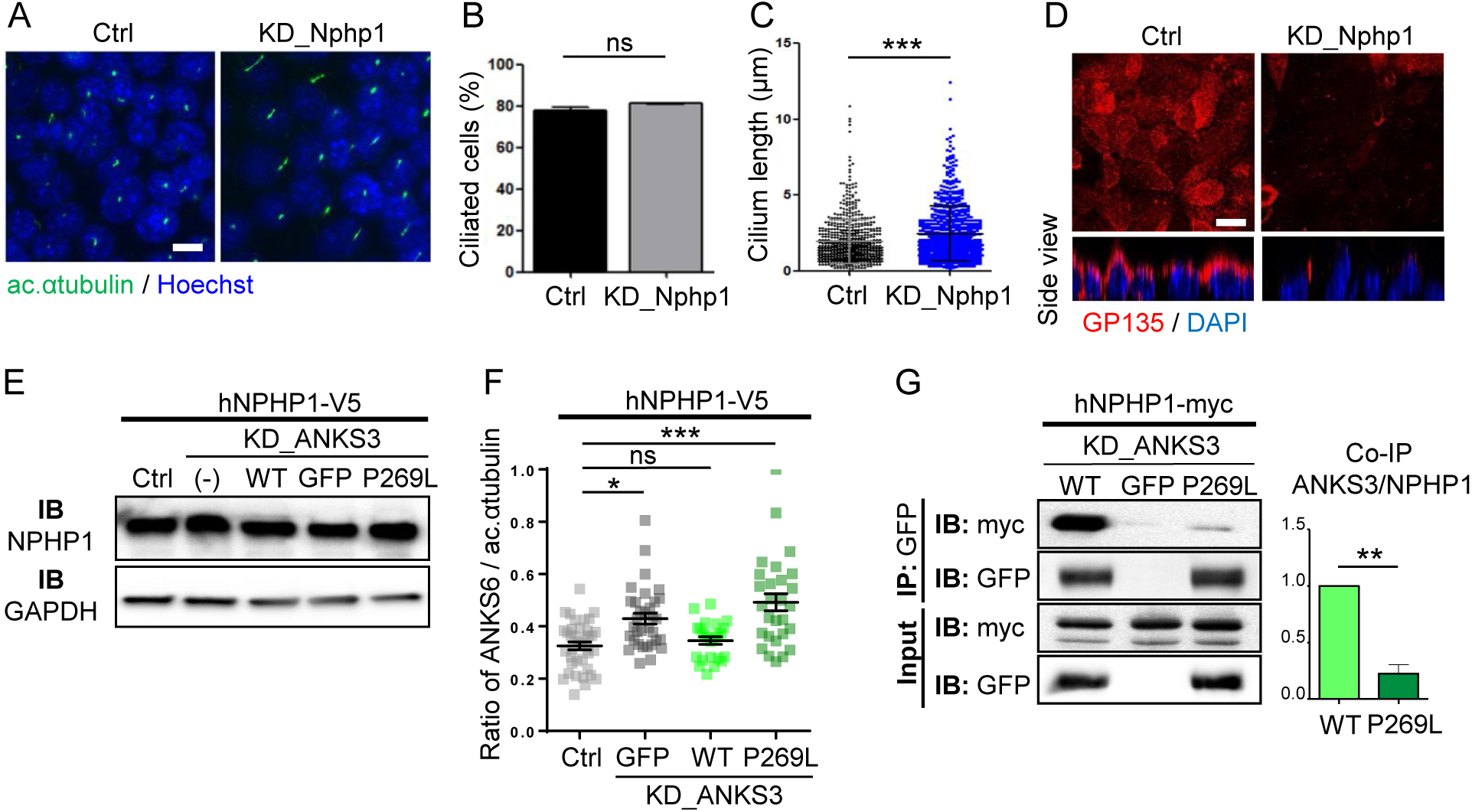
IMCD3 knock-down Nphp1. **(A)** Immunofluorescence of cilia (green, acetylated α-tubulin) in mIMCD3 cells transfected with *Nphp1*-targeted siRNA in IMCD3 cells. Scale bar, 5µm. **(B-C)** Quantification of ciliogenesis (% of ciliated cells, B) and cilium length (C) in control and KD_Nphp1. Values are the mean ± sem. of 3 independent experiments with n >100 cells; *p*-values were determined by Mann-Whitney test. ns, not significant *p* > 0.05, \*\*\**p*≤0.001. **(D)** Subcellular localization of the apical marker GP135 (red) in Control and KD_Nphp1 mIMCD3 cells grown for 6 h in standard medium after Ca^2+^ switch assay. Scale bar, 5µm**. (E)** Western blot of NPHP1 expression in control, KD_GFP, KD_WT, and KD_P269L mIMCD3 cells stably expressing human NPHP1-V5**. (F)** Ciliary distribution of ANKS6 in control, KD_GFP, KD_WT, and KD_P269L mIMCD3 cells stably expressing NPHP1-V5, illustrating the size of the InvsC. Values are the mean ± sem. of 3 independent experiments; *p*-values were determined by one-way ANOVA followed by Dunn’s multiple comparison test; ns: non-significant *p*>0.05, \**p*≤0.05; \*\*\**p*≤0.001. **(G)** Effect of p.P269L mutation on interaction between ANKS3 and hNPHP1-myc. Lysates from mIMCD3 cells transfected with hNPHP1-myc with knockdown of ANKS3 (KD_ANKS3) and re-expressing WT ANKS3-GFP, P269L ANKS3-GFP, or GFP alone were immunoprecipitated with an anti-GFP antibody. Co-immunoprecipitation of GFP-ANKS3 and hNPHP1-myc was quantified by western blot using GFP and myc antibodies. Error bars represent the sem of 3 independent experiments; *p*-values were determined with the Mann-Whitney test, **p≤0.01

**Supplementary Fig. 7.**
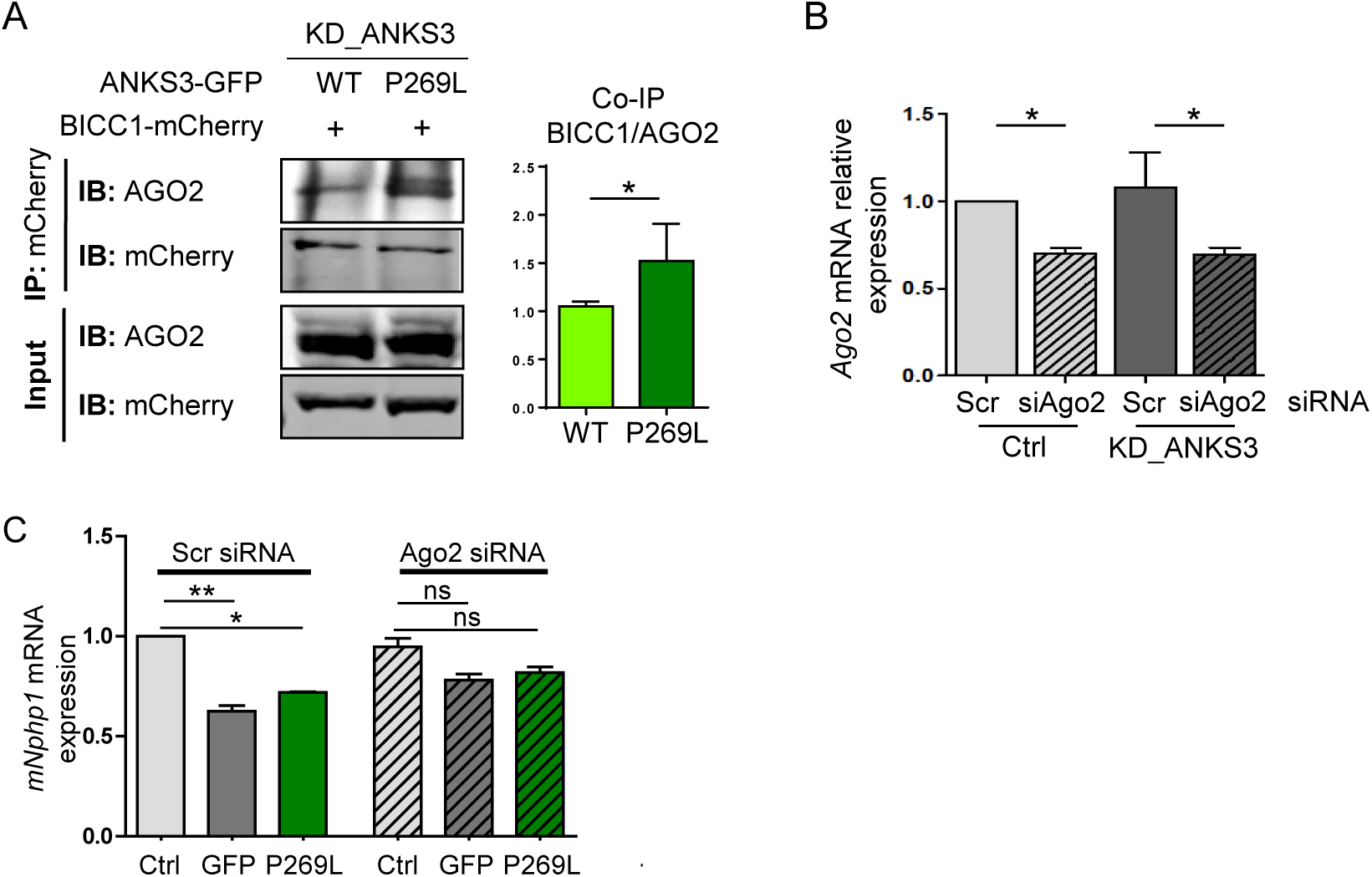
BICC1/AGO2 interaction. **(A)** KD_WT or KD_P269L mIMCD3 cells were transfected with BICC1-mCherry construct and lysates were immunoprecipitated with mCherry antibody. Western blots of immunoprecipitates were performed using mCherry and AGO2 antibodies and interaction of BICC1 with Ago2 was quantified. **(B)** qPCR analysis of relative *Ago2* mRNA expression in mIMCD3 cells transfected with a scramble (Scr) or *Ago2*-targeting siRNA. Values are the mean ± sem. from 3 independent experiments. *p< 0.05 with the Mann-Whitney test. **(C)** qPCR analysis of relative *Nphp1* mRNA expression in Ctrl, KD_GFP and KD_P269L mIMCD3 cells transfected with a scramble siRNA or *Ago2*-targeting siRNA. Values are the mean ± sem. from 3 independent experiments. ns: non-significant p> 0.05, ***p≤0.001 by two-way ANOVA followed by Dunn’s multiple comparison test.

**Table S1:**
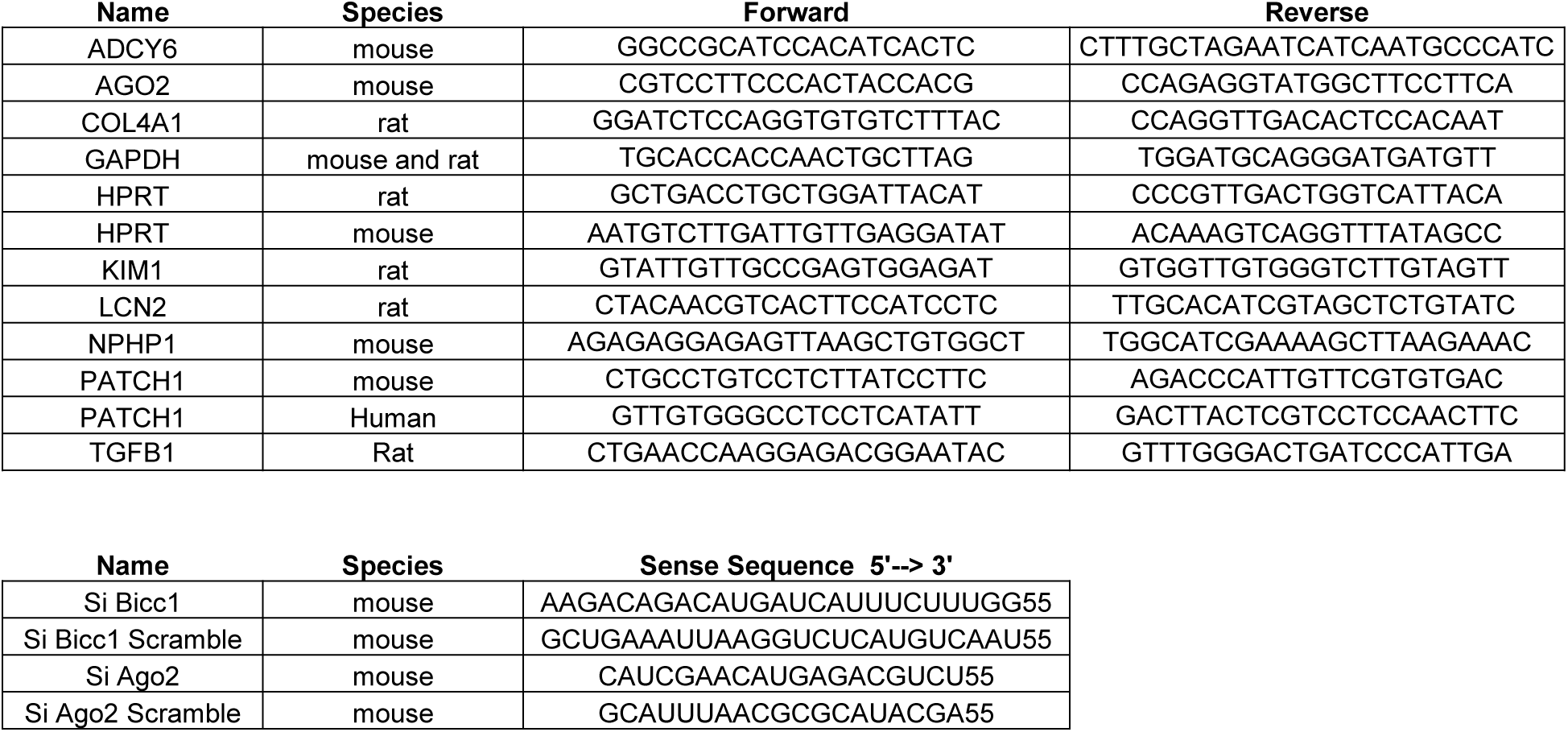
Oligonucleotides.

